# ESRP2-microRNA-122 axis directs the postnatal onset of liver polyploidization and maturation

**DOI:** 10.1101/2024.07.06.602336

**Authors:** Sushant Bangru, Jackie Chen, Nicholas Baker, Diptatanu Das, Ullas V. Chembazhi, Jessica M. Derham, Sandip Chorghade, Waqar Arif, Frances Alencastro, Andrew W. Duncan, Russ P. Carstens, Auinash Kalsotra

## Abstract

Hepatocyte polyploidy and maturity are critical to acquiring specialized liver functions. Multiple intra- and extracellular factors influence ploidy, but how they cooperate temporally to steer liver polyploidization and maturation or how post-transcriptional mechanisms integrate into these paradigms is unknown. Here, we identified an important regulatory hierarchy in which postnatal activation of Epithelial-Splicing-Regulatory-Protein-2 (ESRP2) stimulates biogenesis of liver-specific microRNA (miR-122), thereby facilitating polyploidization, maturation, and functional competence of hepatocytes. By determining transcriptome-wide protein-RNA interactions *in vivo* and integrating them with single-cell and bulk hepatocyte RNA-seq datasets, we delineate an ESRP2-driven RNA processing program that drives sequential replacement of fetal-to-adult transcript isoforms. Specifically, ESRP2 binds the primary miR-122 host gene transcript to promote its processing/biogenesis. Combining constitutive and inducible ESRP2 gain- and loss-of-function mice models with miR-122 rescue experiments, we demonstrate that timed activation of ESRP2 augments miR-122-driven program of cytokinesis failure, ensuring proper onset and extent of hepatocyte polyploidization.

## INTRODUCTION

After fate determination, individual cells within tissues acquire distinct physiological and morphological features as they mature to become fully functional. Maturity arises through a complex interplay of genetic and environmental cues in the perinatal period, which foster cell states that establish/maximize specific functions^1^. Liver maturation is similarly defined through functional specialization, wherein hepatocytes, the primary epithelial cell type of the liver, consolidate a vast array of biosynthetic, metabolic, and detoxification activities after birth, which are zonated based on their location across the lobule^2–4^. These activities involve processing, partitioning, and metabolizing macronutrients; synthesizing bile, glycogen, urea, and serum proteins; and breaking down xenobiotics^5^. The ability of hepatocytes to develop these diverse functional traits is achieved in part by adaptations to postnatal feeding-fasting periods, oxygenation, and hormonal changes that occur during the suckling-to-weaning transition period^6–9^. This also requires precise activation of transcriptional, posttranscriptional, and signaling programs to ensure proper differentiation, growth, and maturation of hepatocytes^10–14^.

Hepatocyte maturity and proliferation are inversely correlated during postnatal development. While hepatocytes are highly proliferative in the fetus, they adapt a quiescent state after birth, undergoing concurrent structural, functional, and metabolic maturation^2, 15^. Notably, this adaptive mature state is reversible, as quiescent hepatocytes can re-enter the cell cycle following injury and regenerate a damaged liver^16–18^. Another defining feature of liver maturation is postnatal polyploidization of hepatocytes. It is estimated that over 70% of rodent and 40% of human hepatocytes have >2n chromosomal DNA content, making liver one of the most polyploid organs in mammals^19, 20^. Polyploid hepatocytes in mice emerge between 2-3 weeks after birth^21^; and the primary mechanism for somatic increase in hepatocyte ploidy is programmed cytokinesis failure, which produces tetraploid and octoploid cells with one or two nuclei^22, 23^. This postnatal surge in ploidy and binucleation overlaps with terminal differentiation and increased size of hepatocytes^24^. It is believed that polyploidization serves as a means to gain specialized functions, boost hepatic gene expression for higher metabolic activity, and provide extra genome copies to buffer against genotoxic damage^25–27^. Several intra- and extracellular factors have been identified that influence hepatic polyploidy, including transcription factors: *E2F1/7/8*^28–30^, and *cMyc*^31^; cell cycle regulators: *p53*^32^, and *Rb*^33^; signaling pathways: *insulin-PI3K-AKT*^34^ and *Hippo-Yap*^35^; and microRNAs: *miR-122*^21^. Yet, how these diverse factors cooperate to balance the polyploidization and proliferation states of maturing hepatocytes is poorly understood.

In this study, we describe a cell type- and developmental stage-specific RNA processing program that guides hepatocyte maturation and polyploidization in murine livers. In-depth molecular and phenotypic analyses identified a critical role for epithelial-specific RNA binding protein ESRP2 in governing hepatocyte size, proliferation, ploidy, and metabolism. Combining single-cell and bulk hepatocyte RNA-seq analyses with targeted deletion of *Esrp2* in mouse livers revealed its requirement in the genesis and maintenance of the adult transcriptome necessary for supporting the mature phenotype of hepatocytes. We also demonstrated that ectopic expression of ESRP2 is sufficient to activate an adult RNA processing program in neonatal livers, evoke premature cell-cycle exit, and induce early maturation and polyploidization of hepatocytes. Next, by CRISPR-mediated FLAG-tagging of ESRP2 allele and using eCLIP to purify native ESRP2 protein-RNA complexes from mice livers, we determined the hepatic ESRP2 RNA-binding landscape at single-nucleotide resolution. Specifically, we discovered robust ESRP2 binding to miR-122 host gene transcript and showed that ESRP2 stimulates the processing and optimal expression of mature miR-122. Finally, by leveraging constitutive/inducible mice models of hepatocyte-specific ESRP2 depletion and overexpression along with miR-122 rescue experiments, we demonstrated that programmed increase in miR-122 processing—via postnatal ESRP2 activation—stimulates cytokinesis failure and expansion of binucleate hepatocytes. Thus, ESRP2 acts in a strict temporal window to activate an RNA processing program, which dictates the terminal differentiation, maturation, and polyploidization of hepatocytes.

## RESULTS

### ESRP2 expression in maturing hepatocytes is regulated transcriptionally and post-transcriptionally

ESRP2 is a conserved RNA binding protein that activates an adult splicing program in hepatocytes^14^; but, how its developmental expression/activity are controlled is undetermined. ESRP2 is absent in fetal hepatocytes but its expression increases gradually during the late gestational period, suggesting a transcriptional regulatory mechanism that modulates *Esrp2* gene expression in a manner which is dependent on developmental cues. The interplay between active and repressive histone modifications often dictates the accessibility of transcriptional machinery to specific gene loci^36, 37^. We therefore analyzed the ENCODE chromatin immunoprecipitation sequencing (ChIP-Seq) data from mice livers and determined the status of chromatin-associated histone modifications and their corresponding marks around the *Esrp2* gene body **(Figure 1A)**. A dramatic shift in the histone marks on *Esrp2* gene locus was evident during the transition from early embryonic stages to adulthood. Particularly, marks of transcriptional activation H3K4me1, H3K4me3, H3K36me3, H3K27ac, and H3K9ac were largely absent over the *Esrp2* gene until birth **(Figure 1A)**. In contrast, the transcriptional silencing mark H3K27me3 was predominant in the embryonic stage but was replaced with activation marks in the adult liver. These data imply that developmental changes in histone modifications facilitate *Esrp2* transcription resulting in its increased mRNA expression.

**Figure 1.**
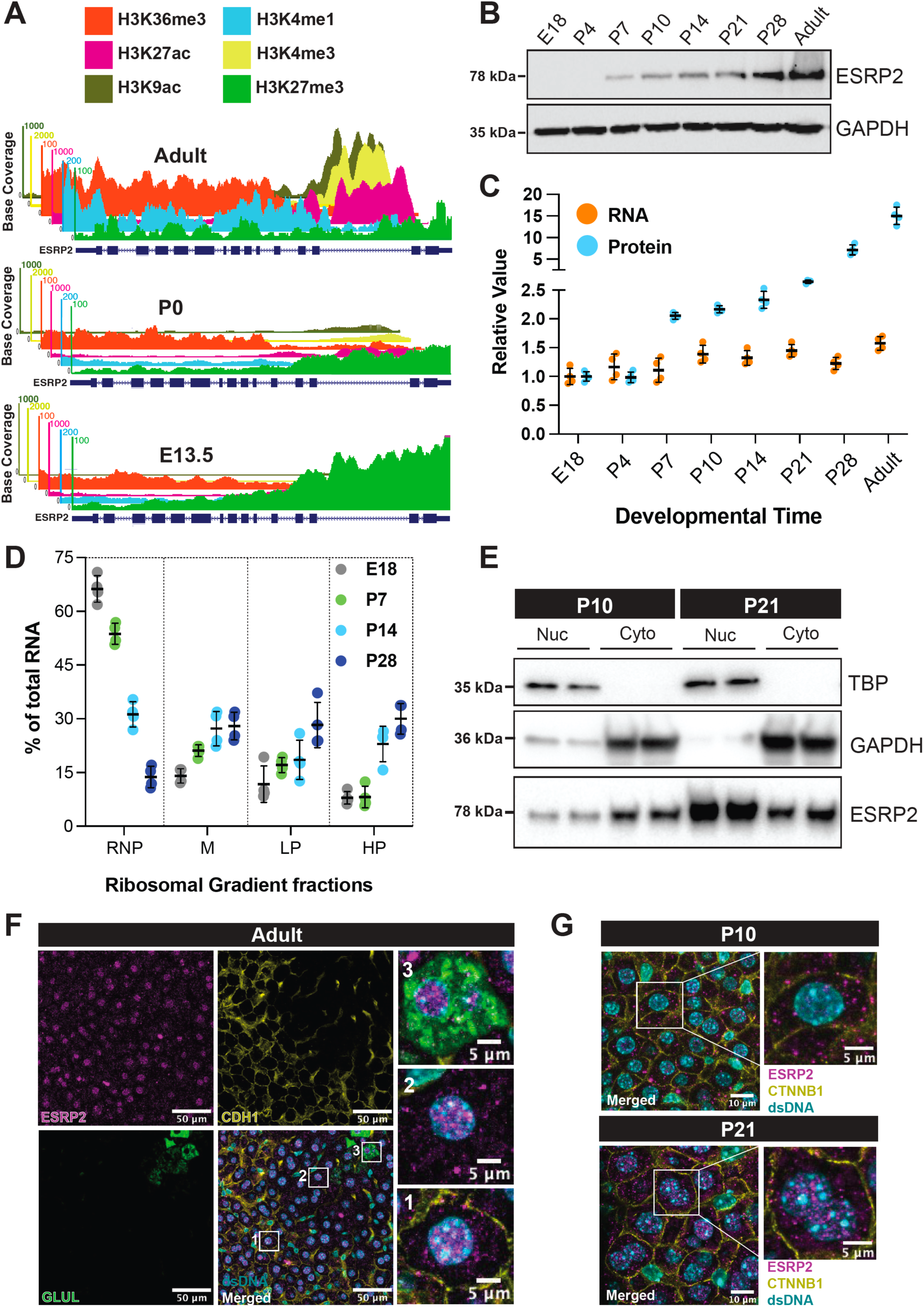
Multi-tiered control of ESRP2 expression during postnatal liver maturation. **(A)** ChIP-seq tracks (from ENCODE) showing changes in histone modifications on *Esrp2* gene locus from Embryonic Day (E) 13.5, Postnatal Day (P) 0 and Adult (P56) mice livers. **(B)** Representative western blot showing changes in ESRP2 protein abundance during postnatal liver development in mice. (n = 4-5 animals, all blots were repeated independently >3 times). **(C)** Relative quantification of *Esrp2* mRNA (qRT-PCR; Taqman probes) normalized to *Gapdh* expression and ESRP2 protein (from panel B) normalized to GAPDH levels during postnatal liver development in mice (n = 4-5). Values are displayed as Mean ± SD. **(D)** qRT-PCR-based quantification of *Esrp2* mRNA in different polysomal fractions of hepatocytes isolated from wildtype mice livers at indicated timepoints. M: monosome, LP: Light polysome, HP: Heavy polysome (n = 3). Values are displayed as mean ± SD. **(E)** Western blots for ESRP2, GAPDH and TBP proteins in nuclear (Nuc) and cytoplasmic (Cyto) fractions from Postnatal Day P10 and P21 mice livers (n = 3). **(F)** Representative Immunofluorescence (IF) images stained for ESRP2 along with periportal (CDH1) and pericentral (GLUL) regions in wildtype adult mouse livers. The boxed 1, 2, and 3 areas represent the pericentral, midlobular and periportal regions. **(G)** IF staining for ESRP2, and a membrane-specific (CTNNB1) marker in wildtype P10 (TOP) and P21 (BOTTOM) mouse livers. dsDNA was labeled with Hoechst 33342 in (F) and (G).

Unexpectedly, while histone modifications changed dramatically from E13.5 to P0, ESRP2 protein expression was only detected starting around P7 **(Figure 1B)**. Further quantification of hepatic ESRP2 mRNA and protein levels exhibited strong upregulation in protein levels (∼8-fold) between P10 and adulthood but showed only modest changes in RNA levels (∼1.5-fold) **(Figure 1C)**. These results suggest that ESRP2 protein abundance in maturing hepatocytes is regulated post-transcriptionally, either at the level of mRNA translation or protein stability (or both). To investigate whether delayed postnatal appearance of ESRP2 was due to increased protein synthesis, we performed polysome profiling of hepatocytes isolated from mice livers^38, 39^ at four developmental time points (E18, P7, P14, and P28). Quantitative RT-PCR (qRT-PCR) analyses of fractionated lysates showed a gradual shift of *Esrp2* mRNA from the mRNP/monosome to polyribosome fractions at P7 **(Figure 1D)**, representing increased accessibility to translational machinery occurring approximately one week after birth.

Given that splicing activity of RNA binding proteins can be coordinated through their nuclear-cytoplasmic distribution, we prepared nuclear and cytoplasmic fractions from hepatocytes isolated from P10 and P21 livers and assayed the relative ESRP2 protein levels in these fractions. The subcellular fractionation results showed that while ESRP2 was primarily cytoplasmic at P10, it became predominantly nuclear by P21 **(Figure 1E)**. Particularly, ESRP2 protein abundance increased sharply in the nuclear fraction at P21, whereas the cytoplasmic levels were unchanged between P10 and P21. We next examined the zonal and subcellular distribution of ESRP2 by immunofluorescence on liver sections that were co-stained with CTNNB1 to mark the cell boundaries or with CDH1 and GLUL to mark the periportal and pericentral hepatocytes, respectively. ESRP2 was uniformly expressed in all hepatocytes across the three zones of the liver lobule **(Figure 1F, Figure S1)**; and similar to the immunoblot results, ESRP2 signal shifted from a predominantly cytoplasmic location at P10 to a predominantly nucleoplasmic location at P21 **(Figure 1G)**. We further noticed that within the nucleoplasm, ESRP2 displayed a punctate staining pattern that concentrated near the euchromatic and away from the heterochromatic centers^40^. Collectively, these data highlight the multi-tier control of ESRP2 expression in maturing hepatocytes, suggesting its stage-specific role in RNA processing during postnatal development.

### ESRP2 is a determinative alternative splicing factor required for complete hepatic polyploidization and maturation

Numerous changes in mRNA abundance and splicing occur within first three weeks after birth^14, 41, 42^, as hematopoiesis and proliferative activity are reduced, and gene products involved in metabolism are induced^3, 43^. During this period, the liver transitions from an immature to a mature state as hepatocytes exit the cell cycle, become polyploid and organize within a zonated lobular structure^4, 20, 44^. Given that these cellular and molecular changes coincide with the postnatal upregulation of ESRP2 and its downstream RNA processing activities **(Figure S2A)**, we set out to investigate ESRP2’s direct contributions towards hepatic maturation using three independent gain- and loss-of-function models. In the first model, the *Esrp2* gene was deleted in the mouse germline, resulting in a constitutive whole-body knockout (KO) **(Figure 2A)**. In the second model, ESRP2 was ectopically expressed in neonatal mice livers **(Figure 2B)**, and for the third model, we generated transgenic mice with *loxP* sites surrounding exon 3-13 of the *Esrp2* gene to acutely disrupt its liver expression using a hepatocyte-specific CRE recombinase **(Figure 2C, Figure S2B)**.

**Figure 2.**
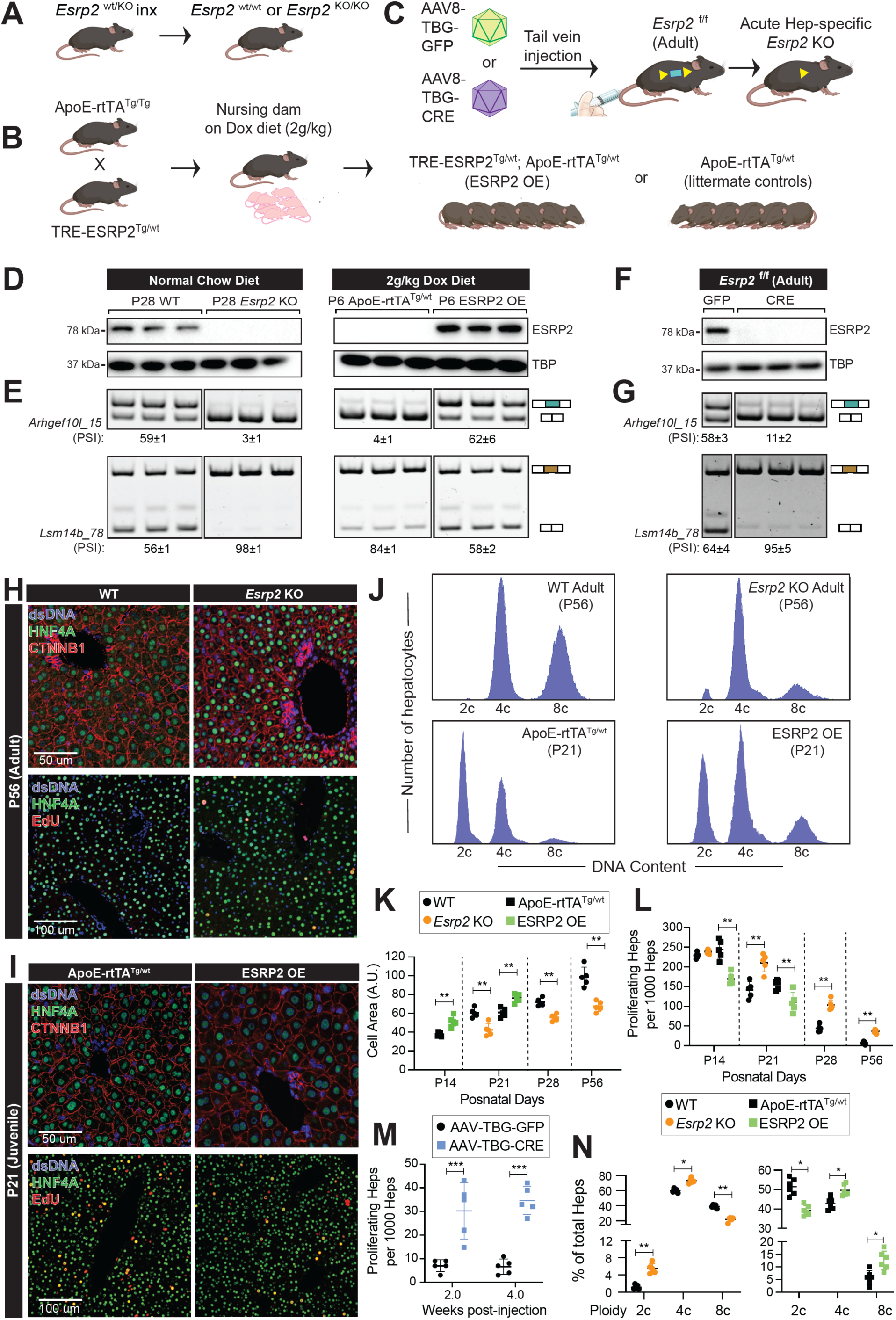
Postnatal onset of ESRP2 activity drives developmental splicing transitions, hepatocyte polyploidization and liver maturation. Respective schematics for generating constitutive and inducible ESRP2 gain- and loss-of-function mice models. **(A)** Germline *Esrp2* deletion mice (*Esrp2* KO), **(B)** Earlier-than-normal doxycycline (Dox)-inducible, hepatocyte-specific, ESRP2 overexpression (TRE-ESRP2 ^Tg/wt^; ApoE-rtTA ^Tg/wt^) in pups, and **(C)** Acute hepatocyte-specific *Esrp2* deletion in adult mice. CRE or GFP transgene expression from AAV8 viruses were driven by the hepatocyte-specific thyroxine binding globulin (TBG) promoter. **(D)** Western blots showing hepatic ESRP2 protein levels, and **(E)** RT-PCR assays showing splicing patterns of ESRP2 target genes in WT, *Esrp2* KO, ESRP2 OE and littermate control mice livers at indicated timepoints. **(F)** Western blots showing hepatic ESRP2 protein levels, and **(G)** RT-PCR assays showing splicing patterns of ESRP2 target genes following acute *Esrp2* deletion (CRE) in adult (P56) livers relative to controls (GFP) at 4 weeks post viral transductions. Representative immunofluorescence (IF) images of livers from **(H)** adult (P56) WT vs. *Esrp2* KO mice, and **(I)** juvenile (P21) ESRP2 OE vs. littermate control mice co-stained for (TOP) CTNNB1, and (BOTTOM) EdU incorporation along with HNF4A (Hepatocyte marker) and DAPI (dsDNA marker). **(J)** Distribution plots for flow cytometry derived ploidy states of hepatocytes from (TOP) adult WT vs. *Esrp2* KO mice and (BOTTOM) juvenile ESRP2 OE vs. littermate control mice. Quantification of hepatocyte **(K)** cell size, and **(L)** proliferation at indicated timepoints from WT, *Esrp2* KO, ESRP2 OE and littermate control mice livers. **(M)** Quantification of hepatocyte proliferation following acute *Esrp2* deletion in adult livers relative to controls at 2 and 4 weeks post viral transductions. **(N)** Quantitative distribution of hepatocytes across ploidy states from (LEFT) adult WT vs. *Esrp2* KO, and (RIGHT) juvenile ESRP2 OE vs. littermate control mice. Data are Mean ± S.D. Two-tailed unpaired T-test with Welch’s correction was used to determine significance between the groups. *p<0.05, **p<0.01, and ****p<0.0001. (n = 4-6 animals/group).

We first determined the spatiotemporal necessity and sufficiency of ESRP2 in coordinating postnatal alternative splicing transitions. Previously, we created a tetracycline-inducible FLAG-tagged ESRP2 transgenic mouse line, which, when crossed with ApoE-rtTA mice, produce TRE-ESRP2; ApoE-rtTA bi-transgenic progenies allowing conditional, doxycycline (Dox)-dependent expression of ESRP2 in hepatocytes^38^. Owing to ESRP2 protein reaching substantial nuclear concentrations only around P14, we induced the bi-transgenic neonates at birth and harvested their liver tissues at P6 as a midpoint between birth and two weeks **(Figure 2B)**. At these early stages of development, the newborn pups depend on the mother’s milk for sustenance. Therefore, we fed the nursing dams Dox-containing diet to dose the pups indirectly through breastfeeding. Western blot analysis confirmed the successful expression of exogenous FLAG-tagged ESRP2 in P6 bi-transgenic livers **(Figure 2D)**. As expected, we did not detect any ESRP2 protein in P6 littermate control or P28 *Esrp2* germline KO livers **(Figure 2D)**. Next, we performed splicing-sensitive RT-PCR assays to interrogate if early induction of ESRP2 promotes a premature advent of the adult RNA splicing program. We chose to evaluate *Arhgef10l* and *Lsm14b* as candidates because both transcripts encode a developmentally regulated alternative exon whose splicing is controlled by ESRP2. For instance, the 15nt alternative exon in *Arhgef10l* was predominantly skipped in P6 but included in P28 livers, whereas the 78nt alternative exon in *Lsm14b* was predominantly included in P6 but skipped more in P28 livers **(Figure 2E)**. The P28 *Esrp2* KOs failed to undergo postnatal transitions and mimicked the wildtype P6 splicing patterns for both transcripts. Remarkably, forced expression of ESRP2 in P6 livers resulted in a neonatal-to-adult switch in splicing for both *Arhgef10l* and *Lsm14b*, demonstrating earlier-than-normal ESRP2 expression is sufficient to induce a premature shift in splicing of target transcripts **(Figure 2E)**.

Because ESRP2 was ablated in the germline, it was unclear whether failure to switch splicing from a neonatal-to-adult pattern resulted from a direct consequence of ESRP2 deficiency or the cascade of events manifested due to absence of ESRP2 during development. To address this, we examined the acute effect of *Esrp2* KO in adult livers by using adeno-associated viral vectors expressing CRE recombinase driven by hepatocyte-specific thyroxine-binding globulin (TBG) promoter **(Figure 2C)**. Robust depletion of ESRP2 protein was evident in the adult *Esrp2* ^f/f^ mice livers two weeks after AAV8-TBG-CRE viral vector transduction compared with AAV8-TBG-GFP controls **(Figure 2F)**. Importantly, acute loss of ESRP2 from adult hepatocytes reverted the splicing of *Arhgef10l* and *Lsm14b* to neonatal patterns—similar to what was observed in the germline KOs **(Figure 2E, 2G)**—indicating continuous expression of ESRP2 in adulthood is necessary to maintain correct splicing patterns of target transcripts. These data provide compelling evidence that ESRP2 is a determinative factor required for both genesis and maintenance of the adult hepatic splicing program, and that its temporally regulated expression ensures timely switch in alternative splicing within maturing hepatocytes.

To determine how the postnatal onset of ESRP2 activity affects liver maturation, we next analyzed hepatocyte size, proliferation, and ploidy content from wildtype, *Esrp2* KO, and overexpression (OE) mice. Liver sections immunostained with HNF4A and CTNNB1 (to mark cell boundaries) showed hepatocytes with significantly reduced cell area and many more hepatocytes per field from P14, P21, P28, and P56 (adult) *Esrp2* germline KO mice compared to those of wildtype controls **(Figure 2H, 2K, Figure S3A)**. We also detected a considerable increase in hepatocyte proliferation—indicated by increased 5-ethynyl-2’-deoxyuridine (EdU) labeling of new DNA synthesis and HNF4A immunostaining of liver sections—from *Esrp2* KO relative to wildtype mice **(Figure 2H, 2L, Figure S3B)**. Importantly, acute depletion of ESRP2 from adult livers led to a significant increase in EdU-labeled hepatocyte population at 2 and 4 weeks after AAV8-TBG-CRE injections compared with AAV8-TBG-GFP controls **(Figure 2M, Figure S4A)**, suggesting increased proliferation resulted directly from ESRP2 deficiency rather than indirectly from a developmental failure to exit the cell cycle. Hepatocyte size in the acute knockouts also trended lower but did not reach statistical significance (**Figure S4B)**. No differences in liver-to-body weight ratios or signs of liver damage were evident at 4 weeks after acute knockout of ESRP2 **(Figure S4C-D).** To further investigate whether premature ESRP2 expression alters the course of hepatic maturation, we induced age-matched TRE-ESRP2; ApoE-rtTA bitransgenic (ESRP2 OE) and littermate control pups with Dox starting at P0 and harvested their livers at P14 or P21. Notably, earlier-than-normal ESRP2 activation evoked premature decrease in hepatocyte proliferation with simultaneous increase in cell size **(Figure 2I, 2K, 2L)**, indicating ESRP2 promotes cell cycle exit and hypertrophic growth of hepatocytes.

Acquisition of polyploidy in the postnatal period is a conserved feature of liver growth and maturation^20^. At birth, majority of mouse hepatocytes are diploid mononucleated cells that undergo normal cycling. However, between P14 and P21, a specialized mitotic program involving programmed cytokinesis failure promotes rapid binucleation and polyploidization of hepatocytes^19, 21, 23^. To determine whether lack of ESRP2 affects ploidy in the liver, we isolated primary hepatocytes from adult wildtype and *Esrp2* KO mice and measured their ploidy states (2c, 4c, and 8c) by flow cytometric quantification of DNA content. As expected, only a tiny fraction of wildtype adult hepatocytes was diploid, and most of the population exhibited tetraploid or octaploid DNA content **(Figure 2J, 2N)**. The *Esrp2* KO livers, however, showed a marked reduction in octaploid hepatocytes, with a concomitant increase in tetraploid and diploid hepatocytes. Consistent with increased EdU labeling, ESRP2 deficiency resulted in a significant increase in the percentage of Ki-67^+^ hepatocytes **(Figure S3B),** indicating reduced ploidy of *Esrp2* KO hepatocytes is not due to their inability to synthesize DNA and/or proliferate. We next evaluated the impact of early ESRP2 induction on the ploidy spectrum of maturing hepatocytes. We therefore administered Dox to TRE-ESRP2; ApoE-rtTA bitransgenic (ESRP2 OE) and littermate control pups starting at P0 and then collected their hepatocytes at P21 to analyze their ploidy states. At P21, only a small percentage of control hepatocytes were octaploid, whereas the majority were diploid or tetraploid **(Figure 2J, 2N)**. However, earlier-than-normal ESRP2 expression boosted the number of octaploid and tetraploid cells in P21 livers **(Figure 2J, 2N)**, indicating premature ESRP2 activation accelerates the postnatal polyploidization of hepatocytes. Together, these gain- and loss-of-function experiments illustrate that ESRP2 is a determinative RNA processing factor that facilitates polyploidization and maturation of hepatocytes.

### Identification of ESRP2-mediated transcriptome changes in maturing hepatocytes

To study mechanisms underlying the immature phenotype of ESRP2-deficient hepatocytes, we performed single-cell RNA sequencing (scRNA-seq) on liver cells isolated from wildtype P14, adult, and *Esrp2* KO mice **(Figure S5A)**. We captured a total of 12,435 cells that were evenly distributed among the three sample groups and had a mean UMI count of 4065 and a median of 1759 genes expressed per cell **(Figure S5B)**. Cell-types were identified based on the top differentially expressed genes and from canonical cell-type-specific markers^41^; which after filtering, dead-cell removal, and batch-correction, yielded 10,026 hepatocytes and 2,409 non-parenchymal cells (NPCs) **(Figure S5C-D)**. Because ESRP2 expression in the liver is hepatocyte-specific, we focused our analyses on hepatocyte populations. As expected, wildtype P14 and adult hepatocytes formed distinct non-overlapping clusters in the UMAP plot, representing stark differences in their transcriptome **(Figure S5E)**. Majority of ESRP2-deficient hepatocytes, however, clustered adjacent to P14 and away from adult hepatocytes.

To determine if all or some of the *Esrp2* KO hepatocytes resemble a P14-like state, we conducted pseudotime ordering of cells through the postnatal maturation process. The majority of P14 and adult hepatocytes resided on the far ends of the pseudotime trajectory; however, ESRP2-deficient hepatocytes occupied a position towards the middle of the maturation trajectory, forming some branches that were not populated by P14 or adult cells **(Figure 3A, B)**. Interestingly, while many ESRP2-deficient hepatocytes ordered closer to the immature P14 hepatocytes, some *Esrp2* KO hepatocytes also ordered near the wildtype adult hepatocytes along the pseudotime. This suggests that despite the lack of ESRP2, a minor proportion of KO hepatocytes may still adapt to mature and support the functions of an adult liver. Next, we assessed the strength of various metabolic pathways between wildtype and *Esrp2* KO hepatocytes. Several mature metabolic gene sets were dampened in *Esrp2* KO hepatocytes, as is evident from a global shift in their pathway score distribution **(Figure 3B, Figure S5F).** Whereas the gene sets belonging to glycolysis, gluconeogenesis, glycogen metabolism, and the pentose phosphate pathway showed the most significant decrease **(Figure 3B)**, other pathways such as fatty acid and amino acid metabolism were only slightly reduced in the *Esrp2* KO hepatocytes **(Figure S5F)**. To explore the gene regulatory networks that might be contributing to the immaturity of ESRP2-deficient hepatocytes, we used the SCENIC pipeline to generate regulons for transcription factors and their co-expressed genes from individual hepatocytes^45^. *Esrp2* KO hepatocytes were particularly stunted in their adult regulons (e.g., HNF4A, DBP, CEBPA), typically activated during postnatal development **(Figure 3C)**. However, regulons that are active in P14 but silenced in adult hepatocytes were less affected by ESRP2 deficiency. This indicates that the lack of ESRP2 hinders the ability of hepatocytes to mature, but these hepatocytes do not regress to a completely immature fetal-like state.

**Figure 3.**
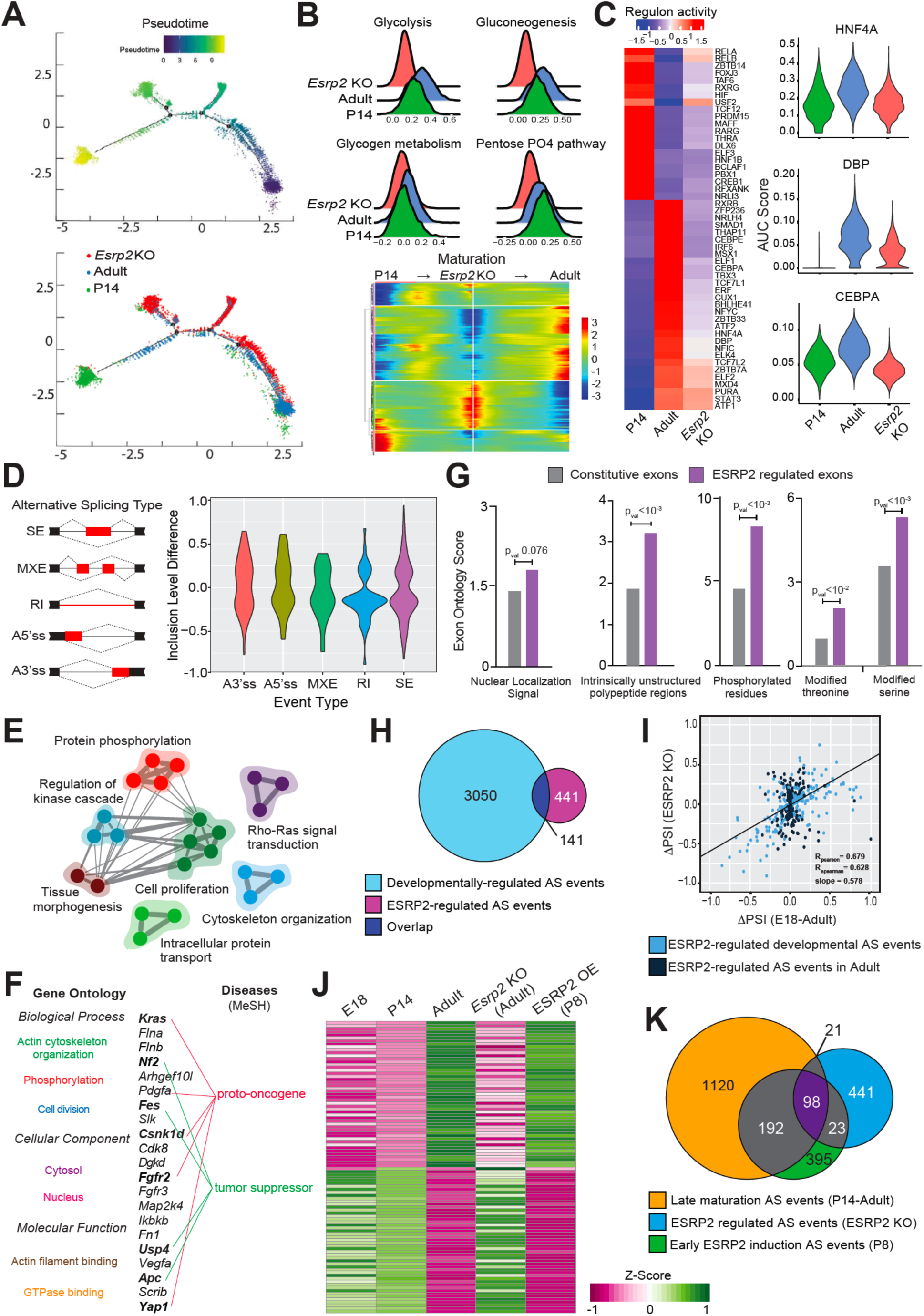
Identification of ESRP2-mediated transcriptome changes in maturing hepatocytes. **(A)** Single-cell RNA-seq analysis showing pseudotime trajectories and beam plots of individual hepatocytes from P14 WT, adult WT, and adult *Esrp2* KO mice livers. Trajectories are colored by pseudotime (top) and sample identity (bottom). Heatmap (right) representing modules of genes that co-vary along the pseudotime in WT maturing vs. *Esrp2* KO hepatocytes. **(B)** Ridge plots showing relative strengths of metabolic pathways in P14 WT, adult WT, and adult *Esrp2* KO hepatocytes. **(C)** SCENIC based analysis of gene regulatory networks in P14 WT, adult WT, and adult *Esrp2* KO hepatocytes. Heatmap (left) depicting regulon activities of transcription factors that show condition-specific variations. Violin plots (right) showing distribution of AUC scores for HNF4A, DBP, and CEBPA regulons across hepatocytes from each condition. **(D)** Violin plots showing distribution of significantly changing alternative splicing (AS) events in adult *Esrp2* KO hepatocytes according to the event types (p<0.05, FDR<0.10, difference in % spliced in [ι1PSI]≥15%, and Junction Counts≥10) obtained from high-resolution bulk hepatocyte RNA-seq of 3 biological replicates from each condition. **(E)** Gene Ontology (GO) network map, and **(F)** Functional categories of significantly enriched pathways and molecular functions for transcripts undergoing differential AS in *Esrp2* KO hepatocytes. **(G)** Exon Ontology based distribution of constitutive and ESRP2-regulated exons according to their encoded protein features. **(H)** Venn diagram, and **(I)** Scatter plot showing overlap of postnatal (P14-Adult) and ESPR2- regulated AS events. **(J)** Row normalized heatmap of Percent Spliced In (PSI) values for AS events regulated in hepatocytes isolated at indicated timepoints from WT, *Esrp2* KO, ESRP2 OE and littermate control mice livers. **(K)** Venn diagram showing intersections of AS events regulated in late phase of hepatocyte maturation, in adult *Esrp2* KO and in earlier-than-normal ESRP2 OE hepatocytes.

Although scRNA-seq provided aggregate information about the differences in hepatocyte cell states between wildtype and *Esrp2* KO livers, it only measured ∼10% of the transcriptome and did not capture any splice variants. Therefore, to assess the global impact of ESRP2 on developmentally regulated splicing, we conducted high-resolution poly(A)^+^ RNA sequencing of wildtype and *Esrp2* KO hepatocytes isolated from 8-week-old mice livers as well E18, P14 hepatocytes from only wildtype livers **(Figure S6A)**. We also investigated the effect of early ESRP2 induction on the splicing pattern of maturing hepatocytes by inducing TRE-ESRP2; ApoE-rtTA bitransgenic and control pups with Dox starting at P0, and then isolating and sequencing their hepatocyte RNAs at P8. ESRP2 deficiency affected the splicing of many more transcripts than abundance, indicating a more significant impact on pre-mRNA processing **(Figure S6B)**. We identified 582 significantly changing splicing events within 394 genes in *Esrp2*-KO compared to WT adult hepatocytes, with an expected distribution of splice event type, and minimal overlap with mRNAs regulated at the abundance levels **(Figure 3D)**.

We noted that the transcripts exhibiting altered splicing patterns in ESRP2-deficient hepatocytes form highly connected networks of biological processes associated with tissue morphogenesis, cell proliferation, and EMT with particular enrichment for molecular functions related to protein phosphorylation, cytoskeletal organization, and Rho/Ras signaling **(Figure 3E)**. Further analysis of the ESRP2-regulated splicing events in hepatocytes identified twenty-one genes encoding components of cell division and/or actin cytoskeletal organization—a subset of which are related to the NCBI MeSH terms: “proto-oncogene” or “tumor suppressor” **(Figure 3F)**. Next, we selected ESRP2-regulated cassette exons with at least 80% sequence conservation between mice and humans (n=462) and probed their protein features and functional properties. Compared to randomized sets of exons, ESRP2-regulated exons showed particular enrichment for nuclear localization signals and intrinsically unstructured polypeptide regions **(Figure 3G)**. We further noticed that the exons containing unstructured regions were significantly enriched for phosphorylation sites, and they often encoded conserved serine/threonine residues that are frequently phosphorylated in response to changes in cellular state and/or activities.

Comparative analysis of the wildtype E18-Adult and ESRP2 KO transcriptomes revealed that almost 1/3^rd^ of differentially spliced mRNAs in ESRP2-deficient hepatocytes revert their splicing pattern towards the neonatal (E18) stage **(Figure 3H and 3I)**. The remaining 2/3^rd^ of mRNAs showed missplicing exclusively in the ESRP2 KO hepatocytes, as their splicing pattern was unaffected during hepatocyte maturation. However, a large but separate set of developmentally regulated events showed only a modest adult-to-neonatal splicing shift in ESRP2 KOs, suggesting other factors might promote these postnatal transitions **(Figure 3H and 3I)**. Thus, our global analysis revealed portions of the hepatocyte transcriptome that undergo dampening of the adult or reactivation of the neonatal splicing program in the absence of ESRP2. Upon analyzing the splicing patterns further, we found that developmentally regulated events can be classified into two classes. Approximately 65% of splicing events undergo “early transitions”, meaning their PSI changes maximally between E18 and P14, whereas the other class of events (∼35%) undergo “late transitions”, meaning their PSI changes maximally between P14 and adult timepoints **(Figure S6C)**. To identify putative regulatory elements (sequence motifs) associated with splicing transitions occurring early or late during hepatocyte maturation, regression analysis between motif counts and splicing changes at each time point was conducted for the enriched hexamers. The first and last 250 nucleotides of the upstream and downstream introns were used in these analyses with the exclusion of the core splice-sites. We identified several motifs in each intronic region, including some that resemble the binding sites of known splicing factors **(Figure S7)**. Amongst them, ESRP binding motif showed a strong association with splicing transitions, particularly in the late postnatal stages. Consistent with the motif analysis and delayed ESRP2 expression during hepatocyte maturation, ESRP2 deficiency mainly affected the late set of splicing transitions. **(Figure 3J, Figure S6D)**. Notably, forced expression of ESRP2 in hepatocytes of P8 livers resulted in a premature neonatal-to-adult shift in splicing of ∼20% of the late events, a subset of which were also regulated reciprocally in the adult ESRP2 KO hepatocytes **(Figure 3K)**.

### Determination of ESRP2 protein-RNA interactions, its binding site preference(s), and direct targets in hepatocytes

To generate the ESRP2-RNA interaction map at a single nucleotide resolution and characterize its hepatic splicing-regulatory-network, we performed eCLIP-seq^46^, a technique in which RNA binding proteins are covalently attached to RNA *in vivo* by UV crosslinking followed by immunoprecipitation, purification, and sequencing. Due to a lack of antibodies to immunoprecipitate ESRP2, we engineered a 2xFLAG tag at the N-terminus of *Esrp2* gene using *CRISPR-Cas9* genome editing in mice **(Figure 4A, B)**. We verified the successful expression of FLAG-tagged ESRP2 in adult transgenic mice livers without any noticeable phenotypic differences, including ESRP2 protein levels or changes in PSI of target exons when compared to wild type livers **(Figure 4C)**. Freshly isolated primary hepatocytes from FLAG-ESRP2 transgenics were UV-exposed and protein-RNA complexes immunoprecipitated with anti-FLAG antibody **(Figure 4D).** We isolated and sequenced 50-200 nucleotide RNA fragments that were cross-linked to and co-purified with FLAG-ESRP2. After mapping and filtering, reads from biological replicates were merged to determine the distribution of ESRP2 binding sites by clustering the overlapping reads and identifying statistically significant peaks relative to Input. FLAG-ESRP2 eCLIP tags were broadly distributed across the transcriptome, with 90% of peaks mapping to 685 annotated RefSeq genes and the majority (∼85%) residing within intronic (35.9%) and 3’-UTR (50.8%) elements **(Figure 4E**), suggesting that at least 7% of mouse genes are subject to ESRP2 regulation in hepatocytes.

**Figure 4.**
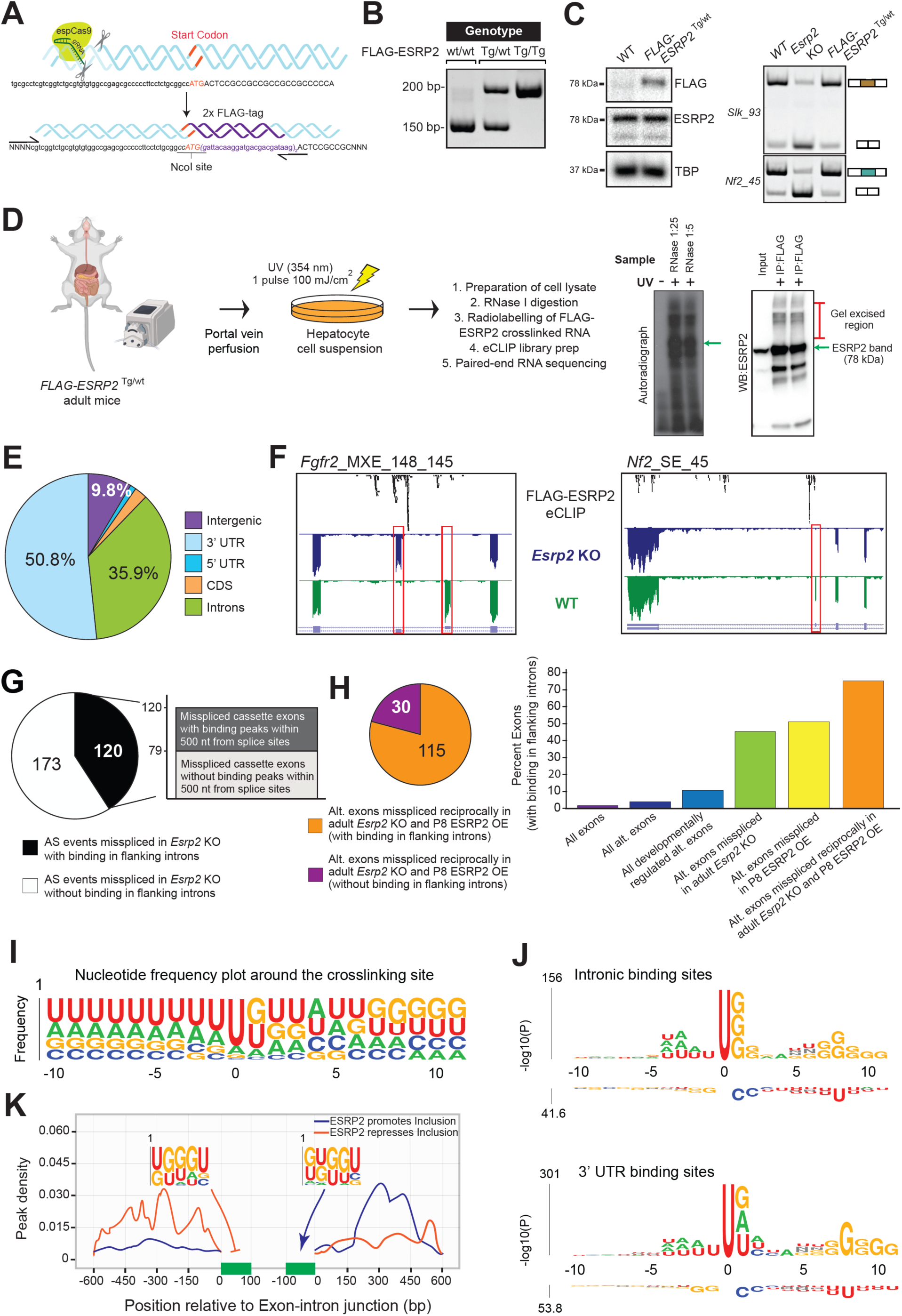
Determination of ESRP2 protein-RNA interactions, binding site preference(s), and direct targets in hepatocytes. **(A)** Schematic representation of CRISPR/Cas9 approach to insert a 51nt DNA fragment encoding 2x FLAG tag downstream of ESRP2 start codon in the mouse genome. Relevant genomic sequence with guide RNA location is shown. **(B)** Genotyping results showing successful insertion of 2x FLAG tag in the ESRP2 N-terminus using primers that amplify products for FLAG-ESRP2 ^wt/wt^ (wildtype), FLAG-ESRP2 ^Tg/wt^ (hemizygous) and FLAG-ESRP2 ^Tg/Tg^ (homozygous) transgenes in mice. **(C)** Western blots (LEFT) showing expression of FLAG-ESRP2 protein, and RT-PCR assays (RIGHT) showing splicing patterns of ESRP2 target transcripts *Slk* and *Nf2* in WT, *Esrp2* KO and FLAG-ESRP2 ^Tg/wt^ mice. **(D)** Strategy for pulldown of hepatic ESRP2 bound RNA and eCLIP analysis from FLAG-ESPR2 ^Tg/wt^ mice. Representative autoradiograph and western blots indicating successful pulldown of FLAG- ESRP2 protein and its crosslinked RNA is shown on the right. **(E)** Transcriptome-wide distribution of ESRP2 binding sites across various genic elements **(F)** Representative genome browser tracks for ESRP2-regulated exons (boxed in red) with FLAG-ESRP2 eCLIP tags (black) in the surrounding intronic regions. Corresponding RNA-seq tracks from WT (green) and *Esrp2* KO (blue) hepatocytes are shown below. Pie charts showing breakdown of **(G)** All ESRP2-regulated alternative exons, and **(H)** Alternative exons reciprocally regulated in adult *Esrp2* KO and P8 ESRP2 OE hepatocytes with and without ESRP2 binding in the flanking introns. **(H, right)** Bar plot showing percentages of alternative exons with eCLIP tags in flanking introns for various subsets. **(I)** Nucleotide Logo Plot showing frequency of the four ribonucleotides at and around the crosslink sites identified from CIMS. **(J)** Top *k*-mers (1-4 nt) enriched and de-enriched at +/- 10 nucleotide positions around crosslink sites identified within introns (TOP) and 3’-UTRs (BOTTOM). **(K)** Density plot of ESRP2 binding peaks near alternative exons whose inclusion is promoted (Blue) or repressed (orange) by ESRP2. The 600 nt region of upstream and downstream flanking introns is shown. ESRP2 preferred binding motifs for Included and Repressed exons are also indicated.

We next integrated the *Esrp2* KO RNA-seq and FLAG-ESRP2 eCLIP-seq datasets to identify pre-mRNAs having eCLIP-tags within close proximity of alternatively spliced exons, which were also misspliced in ESRP2-deficient hepatocytes. We detected robust binding clusters within 500 nucleotides upstream and downstream of exons whose PSI values were significantly altered in *Esrp2* KO hepatocytes **(Figure 4F, Figure S8).** Notably, the global intersection of two datasets revealed that only 1/3^rd^ of variably spliced events in *Esrp2* KOs had a binding peak nearby, which improved to 50% if we focused on cassette exons **(Figure 4G),** suggesting ESRP2 regulates a broad splicing program in hepatocytes via series of primary (*direct*) and secondary (*indirect*) targets. To further distinguish direct from indirect splicing targets, we compared ESRP2 binding preferences across constitutive and alternative exons expressed within maturing hepatocytes. Approximately 2% of “All exons” and 5% of “All alternative exons” had ESRP2 binding in flanking introns; however, this percentage increased to ∼10% if the alternative exons were also developmentally regulated **(Figure 4H)**. ESRP2- regulated exons, which exhibited significant PSI change in ESRP2 depleted or overexpressing hepatocytes, showed noticeably higher binding proportions (∼50%) in surrounding introns. Of note, exons reciprocally regulated in *Esrp2* KO and OE datasets had the highest percentage of binding (∼80%), providing a strong likelihood for direct regulation by ESRP2 **(Figure 4H)**.

To investigate the sequence features of ESRP2-binding sites, we employed an independent read mapping and analysis pipeline that leverages cross-link-induced mutation sites (CIMS) and accurately maps positions of protein-RNA cross-linking in CLIP data^47^. Uridine was the most preferred nucleotide at the crosslink site and was also enriched at the flanking positions **(Figure 4I)**. Guanine and adenine were the second and third most common nucleotides at other positions. We next evaluated nucleotide frequencies in the sequences surrounding ESRP2 crosslink sites (−10 nt to +10 nt) separately within introns and 3’ UTRs **(Figure 4J)**. The *de novo* k-mer enrichment analysis identified UUG- and UGG-containing motifs as the top enriched sequence elements with cross-linking occurring at U residues. Similar UG-containing sequence motifs for ESRP proteins have been identified *in vitro* by SELEX-seq^48^ and RNA bind-n-seq^49^, confirming the accuracy of our *in vivo* ESRP2-binding site determination with FLAG-ESRP2 eCLIP.

Previous studies have shown that linear sequence motifs alone are often insufficient to fully capture the binding specificities of RBPs; and other contextual features such as RNA secondary structure and surrounding base composition contribute to binding specificity^49, 50^. Proteins with a single RNA binding domain (RBD) are believed to interact with 3-5 contiguous RNA bases, whereas proteins with multiple RBDs can interact with bipartite motifs separated by one or more bases, where the two cores in bipartite motifs are bound by distinct RBDs within the same protein^51, 52^. We found that ESRP2, which contains three RRM domains^53^, prefers binding UG dinucleotides followed by a Poly G-tract spaced by six nucleotides, with no preference for specific bases in the intervening spacer **(Figure 4J)**. These results raise a possibility where the downstream Poly G sequence might promote or stabilize ESRP2 interactions with the primary UG-rich motif, and thereby increase its RNA binding specificity.

Finally, to characterize the positional effects of ESRP2 protein-RNA interactions on splicing outcomes at a transcriptome-wide scale, we plotted the density of eCLIP peaks within 600 nucleotides upstream and downstream of cassette exons whose inclusion was repressed (Red) or promoted (Blue) by ESRP2 **(Figure 4K)**. This analysis revealed that ESRP2 binding to the intron downstream of a regulated exon was generally associated with ESRP2 mediated promotion of exon inclusion, whereas binding to the upstream intron was associated with exon skipping. Similar position-dependent RNA regulatory maps have been reported for other splicing regulators, which function in a cell-type or condition-specific manner^54^. Collectively, our integrated RNA-seq and eCLIP-seq analyses established that position-dependent ESRP2 binding facilitates the postnatal splicing switch for a functionally coherent set of RNAs, which support physiological specialization and maturation of hepatocytes.

### ESRP2 binds miR-122 host gene transcript and promotes its processing/biogenesis during postnatal liver development

Because programmed cytokinesis failure drives hepatocyte binucleation and polyploidy^20^, we reasoned that ESRP2-mediated post-transcriptional processing of one or more cytokinesis effector(s) might directly contribute towards postnatal polyploidization of hepatocytes. Prior studies have demonstrated that liver-enriched microRNA, miR-122, is induced during postnatal development, and it stimulates physiological polyploidization of hepatocytes by inhibiting the expression of a set of pro-cytokinesis targets^21^. Surprisingly, in our mapped FLAG-ESRP2 eCLIP-seq reads, we detected extensive ESRP2 binding to the miR-122 host gene transcript **(Figure 5A).** Moreover, while the expression of miR-122 nascent transcript was significantly increased, mature miR-122 levels were reduced in ESRP2-deficient hepatocytes compared to wildtype controls, pointing towards a processing defect **(Figure 5A).** To further validate this observation, we employed a Taqman qRT-PCR assay to compare mature miR-122 levels in wildtype and ESRP2-deficient livers across six postnatal timepoints (P0, P7, P14, P21, P28, and P56). In agreement with earlier reports^21, 55^, hepatic miR-122 expression surged during the first four weeks after birth **(Figure 5B)**. However, starting at P14 and continuing through adulthood, ESRP2 KO livers exhibited consistently lower mature miR- 122 levels relative to wildtype counterparts **(Figure 5B)**.

**Figure 5.**
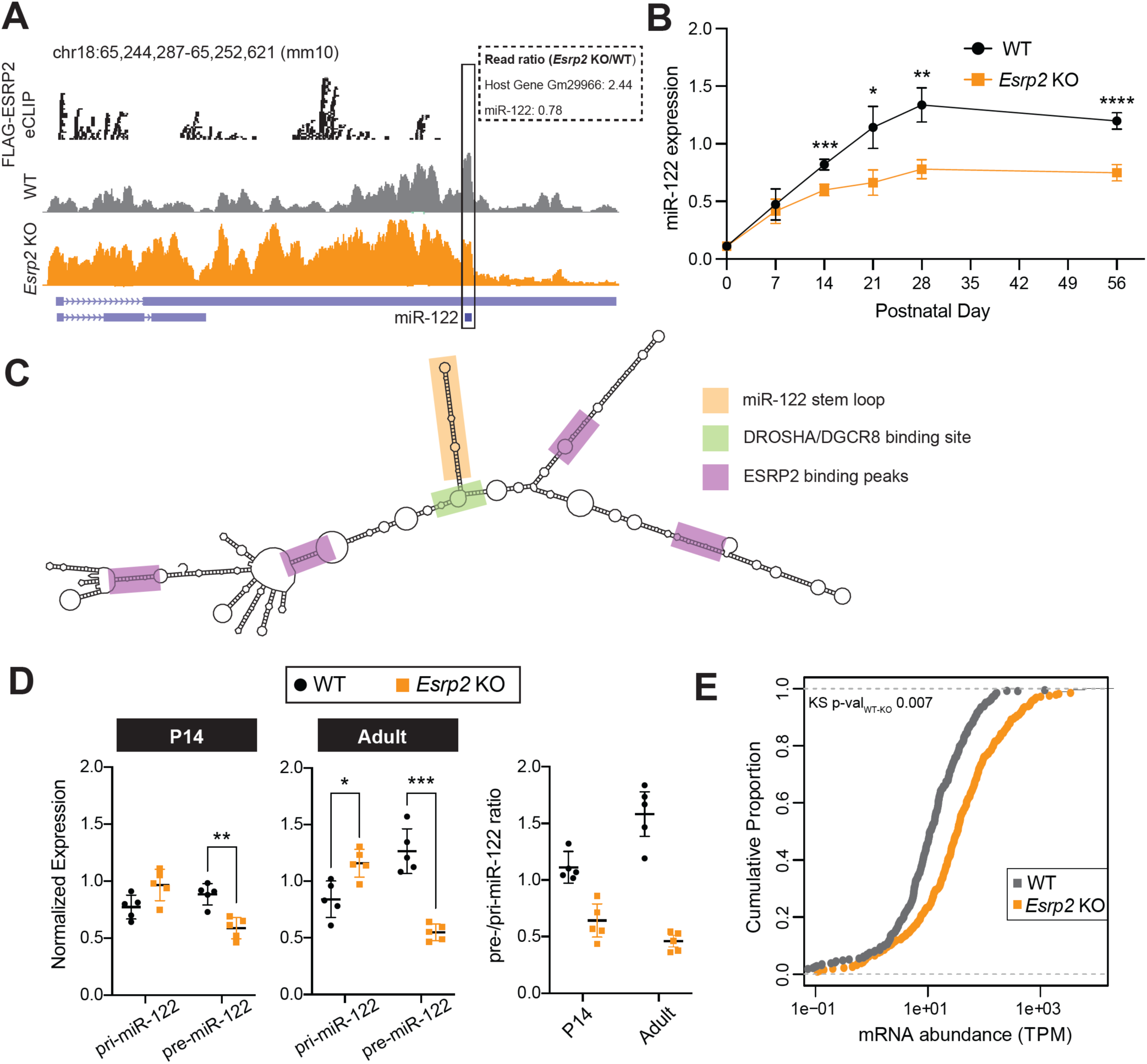
ESRP2 binds primary miR-122 host gene transcript and stimulates its processing/biogenesis in maturing hepatocytes. **(A)** Genome browser view of primary miR-122 host gene Gm29966 with ESRP2 eCLIP tags (black) with corresponding RNA-seq tracks from WT (grey) and *Esrp2* KO (orange) hepatocytes. **(B)** qRT-PCR derived mature miR-122 expression levels at indicated timepoints from WT and *Esrp2* KO mice livers. Data is normalized relative to U6 snRNA. Data are Mean ± S.D (n=5 animals/group). Two-way ANOVA (corrected for multiple comparisons) was used to determine significance between the groups; *p<0.05, **p<0.01, ***p<0.001 and ****p<0.0001. **(C)** RNAfold predicted structure of the +/- 1000 nt region around the miR122 hairpin within the primary transcript. miR-122 hairpin, DROSHA/DGCR8 and ESRP2 binding sites are highlighted. **(D)** qRT-PCR derived pri-miR-122 and pre-miR-122 transcript levels in P14 and Adult (P56) livers from WT and *Esrp2* KO mice. Relative ratios of pre-/pri-miR-122 transcripts are shown on the right. Data are Mean ± S.D (n = 5 animals/group). Two-way ANOVA (corrected for multiple comparisons) was used to determine significance between the groups. *p<0.05, **p<0.01, and ***p<0.001. **(E)** Cumulative plot of mRNA abundance for miR-122-regulated transcripts in RNA-seq data from WT and *Esrp2* KO hepatocytes. TPM; transcripts per million. p-value was calculated using Kolmogorov-Smirnov test.

The canonical microRNA biogenesis pathway involves two sequential processing events catalyzed by RNase III enzymes. In the nucleus, the microprocessor complex involving Drosha/DGCR8 cleaves primary microRNA transcripts (pri-miRs) to produce precursor microRNAs (pre-miRs), which are exported to the cytoplasm where Dicer processes them into mature microRNAs that are eventually loaded into the RISC complex^56, 57^. Upon further analysis of eCLIP data, we noted four distinct clusters of ESRP2 cross-linked fragments that mapped to the UG-rich segments of miR-122 host transcript both upstream and downstream of the stem-loop region **(Figure 5A)**. These ESRP2 binding clusters (>30 reads) were adjacent to the Drosha/DGCR8 cleavage site within the RNAfold^58^ predicted secondary structure and likely facilitate binding to the base of the miR-122 stem loop **(Figure 5C)**. To determine which step of miR-122 biogenesis was affected by ESRP2, i.e., whether ESRP2 facilitates Drosha/DGCR8- mediated processing of the *nascent* or Dicer-mediated processing of the *precursor* miR-122 transcript, we quantified both pri-miR-122 and pre-miR-122 transcript levels in P14 and adult wildtype as well as ESRP2 KO mice livers. Notably, ESRP2 deletion resulted in significant accumulation of unprocessed pri-miR-122 transcripts, with a concomitant reduction in the levels of processed pre-miR-122 transcripts **(Figure 5D)**. Based on the disparate expression patterns of pri- and pre-miR-122 in ESRP2-deficient livers, we conclude that nuclear activity of ESRP2 boosts miR-122 biogenesis by promoting Drosha-mediated cleavage. This conclusion is further supported by the fact that miR-122 levels are closely associated with ESRP2 protein expression during a narrow window of postnatal liver development. For instance, between P0 and P14, miR-122 levels increase by nearly ten-fold **(Figure 5B)**, while ESRP2 protein abundance increases by sixteen-fold **(Figure 1C)**. Also, significant reduction in pre- and mature miR-122 transcripts in ESRP2-deficient hepatocytes is evident only after P14, same timeframe when ESRP2 protein levels rise naturally and its localization becomes progressively nuclear **(Figures 1C, 1D and 5B)**.

miR-122 antagonizes the expression of a large proportion of hepatic mRNAs that encode for a variety of developmental, metabolic, and homeostatic functions^59^. We, therefore, posited that reduced miR-122 levels in ESRP2-deficient hepatocytes may result in chronic de-repression of crucial targets and pathways. Accordingly, we queried the expression of direct miR-122 target gene set in ESRP2 KO hepatocytes, which was derived by intersecting previously published Ago-CLIP-seq and RNA-seq datasets from miR-122 KO and control mice livers^59^. Our comparison of wildtype and ESRP2 KO transcriptomes indicated that in the absence of ESRP2, abundance of miR-122 target mRNAs increases significantly relative to non-targets **(Figure 5E, S9A)**. This was particularly evident for transcripts encoding proteins involved in cell-cell junctions, cytoskeletal rearrangement, and cell division **(Figure S9B).** Collectively, these results support a model where ESRP2 binds multiple UG-rich segments within pri-miR-122 host transcript to stimulate DROSHA-mediated post-transcriptional processing of miR-122.

### Timed activation of ESRP2-miR-122 axis promotes hepatic polyploidization

The data presented above revealed an ESRP2-miR-122 axis that is temporally activated in hepatocytes approximately two weeks after birth. To assess the physiological contribution of this axis, we asked whether ESRP2-driven miR-122 biogenesis potentiates postnatal hepatic binucleation and polyploidization. We first confirmed if ESPR2 was able to increase miR-122 levels *in vivo*, for which we induced TRE-ESRP2; ApoE-rtTA bitransgenic pups with Dox starting at P0 and harvested their livers at P14 and P21 **(Figure 2, 6A)**. Of note, ectopic expression of ESRP2 led to an increase in miR-122 levels in both P14 and P21 livers **(Figure 6B)**. We also measured the mRNA levels of cytokinesis-associated factors, *Rhoa*, *Mapre1*, *Racgap1*, and *Slc25a34*, which are direct targets of miR-122 and, when depleted forcibly, cause a marked increase in binucleate hepatocytes^21^. We observed a significant downregulation of all four factors in ESRP2 overexpressing livers in comparison to uninduced littermate controls **(Figure 6C)**. These data illustrate that earlier-than-normal ESRP2 activation augments the miR-122-driven program of cytokinesis failure and thereby accelerates the ploidy spectrum of hepatocytes.

**Figure 6.**
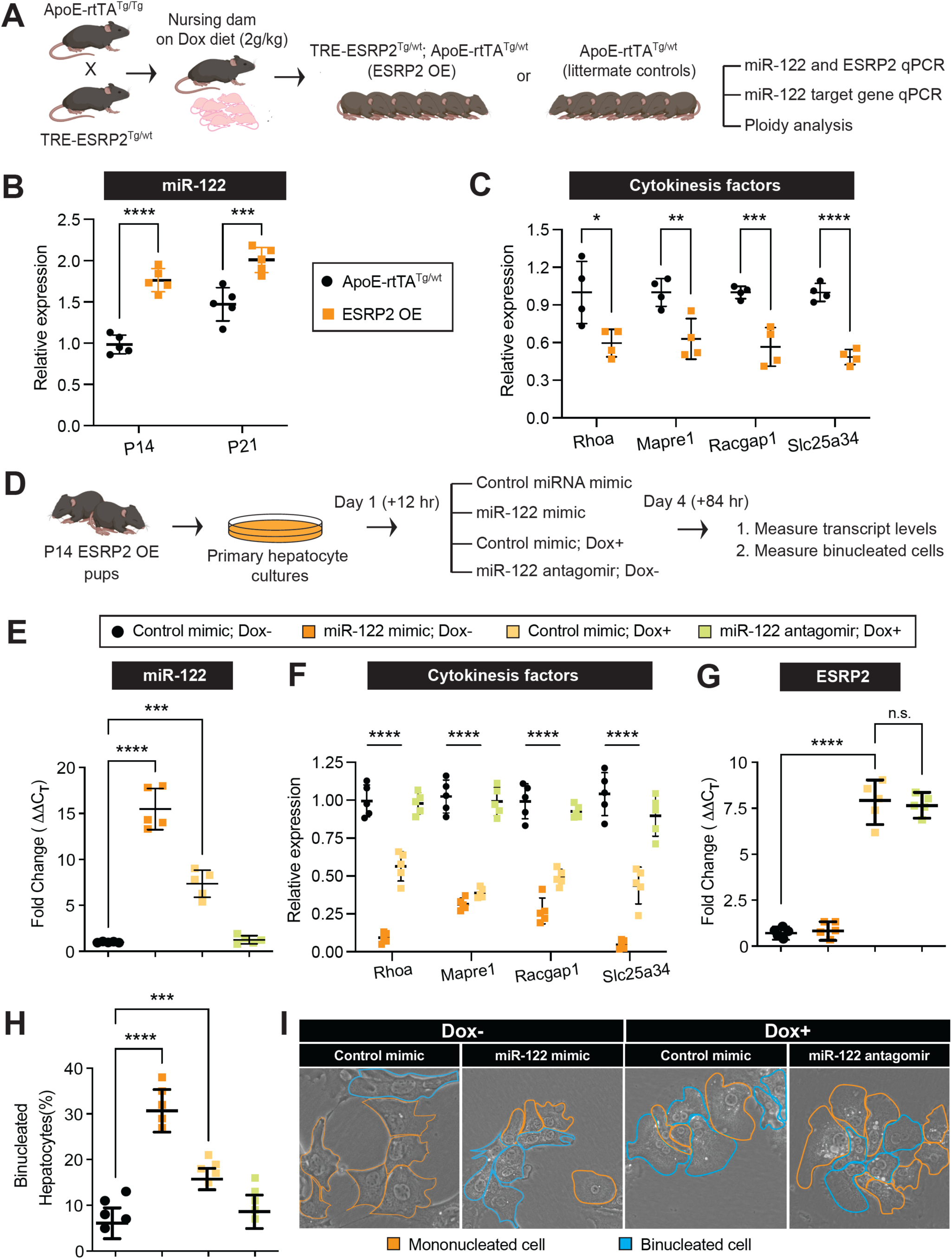
Timed activation of ESRP2-miR-122 axis promotes hepatocyte binucleation. **(A)** Experimental schematic to assess the temporal role of ESRP2 in regulating hepatic miR-122 biogenesis and its target transcript expression following earlier-than-normal ESRP2 OE in pups. Relative levels of **(B)** mature miR-122 (qRT-PCR; Taqman probes) normalized to U6 snRNA, and **(C)** miR-122 target transcripts (qRT-PCR; Sybr green assays) normalized to *Gapdh* in ESRP2 OE and control pup livers at indicated postnatal timepoints. Values are displayed as Mean ± SD (n=5 mice/group). Two-tailed unpaired T-test with Welch’s correction was used to determine significance between the groups. *p<0.05, **p<0.01, ***p<0.001, and ****p<0.0001. **(D)** Experimental schematic of miR-122 inhibition in control and ESRP2 OE primary hepatocytes isolated from P14 ESRP2 OE pup livers and assayed at 4 days post-treatment(s). Relative levels of **(E)** mature miR-122 (qRT-PCR; Taqman probes) normalized to U6 snRNA, **(F)** miR-122 target transcripts (qRT-PCR; Sybr green assays), and **(G)** ESRP2 normalized to *Gapdh* across indicated conditions in (D). **(H)** Percentage of binucleate hepatocytes across indicated conditions in (D), and **(I)** Representative images of mono and binucleate hepatocytes across indicated conditions in (D). Values in (E), (F), (G), and (H) are displayed as Mean ± S.D. Two-way ANOVA (corrected for multiple comparisons) was used to determine significance between the groups. Each data point represents a biological replicate for which cells were derived from a single P14 ESRP2 OE pup. All data are representative of two independent trials. *p<0.05, **p<0.01, ***p<0.001, and ****p<0.0001.

To further demonstrate that the newly elucidated ESRP2-miR-122 axis critically contributed to the postnatal polyploidization of hepatocytes, we performed a functional rescue experiment by breaking the pathway with miR-122 antagomir. For this purpose, we first prepared primary hepatocyte cultures from P14 TRE-ESRP2; ApoE-rtTA bitransgenic mice livers and exposed them to either vehicle or Dox-containing mitogenic growth media **(Figure 6D)**. Over 95% of mouse hepatocytes at P14 stage are diploid, and their binucleation status can be tracked over time *in vitro*^21^. Accordingly, we co-transfected cells with control, miR-122 mimic or miR-122 antagomir oligonucleotides in the presence or absence of Dox. Four days later, cells were either fixed and processed for binucleation analysis or lysed to extract total RNA to measure the transcript levels of ESRP2, miR-122 and specific cytokinesis factors **(Figure 6D)**. As anticipated, miR-122 mimic suppressed the expression of cytokinesis factors: *Rhoa*, *Mapre1*, *Racgap1*, and *Slc25a34* **(Figure 6E and F)**, and miR-122-supplemented hepatocytes were enriched for binucleate cells when compared with scrambled control **(Figure 6H, I)**. Importantly, similar to the *in vivo* mice studies, transient induction of ESRP2 in primary cultures was sufficient to increase miR-122 levels, evoke reduction in *Rhoa*, *Mapre1*, *Racgap1*, and *Slc25a34* mRNAs, and stimulate hepatocyte binucleation **(Figure 6E, F, and H)**. Finally, to verify that increased binucleation upon ESRP2 induction was mediated through miR-122, we co-transfected miR-122 antagomir in Dox-induced TRE-ESRP2; ApoE-rtTA bitransgenic hepatocyte cultures. Supplementing ESRP2-expressing hepatocytes with miR-122 antagomir reduced not only miR-122 levels but also restored the expression of cytokinesis factors and reversed ESRP2-driven accumulation of binucleate cells **(Figure 6E, F and H).** ESRP2 expression itself, however, remained unchanged in the antagomir-treated cells **(Figure 6G).** These data demonstrate that miR-122 upregulation critically contributes to the ESRP2-driven increase in hepatocyte binucleation and polyploidy.

## DISCUSSION

The postnatal period of mammalian development is important as numerous changes take place in the liver at this time^2, 6, 15^. The suckling-to-weaning transition period (P14-P21 in rodents) is particularly significant for liver maturation as hepatocyte size, ploidy, and metabolism increase markedly during this timeframe^23^. Yet, the driving factors of liver maturity, such as how hepatocytes exit the cell cycle, become polyploid, or acquire specialized functions are inadequately understood. Based on our findings, we posit a model of posttranscriptional regulatory hierarchy for liver maturation and polyploidization wherein a programmed increase in miR-122 processing—via postnatal ESRP2 activation—promotes cytokinesis failure and supports diploid-to-polyploid conversion of hepatocytes. *Five* principal observations support this model.

*First*, the normal emergence of polyploid hepatocytes coincides with ESRP2 expression and its downstream RNA processing activities. *Second*, both germline and acute depletion of ESRP2 from mice livers result in reduced hepatic maturity—indicated by increased proliferation, decreased size, and reduced ploidy spectrum of ESRP2-deficient hepatocytes. *Third*, earlier-than-normal activation of ESRP2 in neonatal livers impedes proliferation, evokes cellular hypertrophy, and accelerates hepatocyte maturity and polyploidy. *Fourth*, ESRP2 is required for the *genesis* and *maintenance* of the adult liver transcriptome, including processing and optimal expression of mature miR-122. *Fifth*, timed processing and biogenesis of miR-122 by ESRP2 represses cytokinesis-associated factors, promoting cytokinesis failure and postnatal expansion of binucleate hepatocytes. ESRP2 deficiency derepresses these factors, resulting in deficits in the temporal specification and complete polyploidization of hepatocytes.

RNA processing and miRNA-mediated gene silencing are fundamental mechanisms that fine-tune gene expression to regulate tissue development and physiology^60–63^; and their dysregulation is associated with an increasing number of human diseases, including cancer^64–66^. The interplay between miRNAs and RBPs leads to global changes in gene expression wherein miRNA(s) control expression of RBPs to regulate tissue-specific RNA processing decisions^67–71^ or RBPs modulate miRNA biogenesis, stability, and localization through alternative processing and/or maturation^72–78^. However, most studies interrogating relationships between miRNAs and RBPs utilize *in vitro* approaches or cell culture models. Here, we identified the post-transcriptional ESRP2-miR-122 axis operative *in vivo* during a critical period of postnatal liver development that enables functional maturation and polyploidization of hepatocytes.

ESRPs are a conserved family of epithelial-specific RBPs that exert broad control over many physiological processes, including tissue regeneration and the epithelial-mesenchymal transition (EMT) during development and cancer progression^53, 79–81^. Most epithelial tissues express both ESRP1 and 2, but the liver is unique as ESRP2 is the sole paralogue present in human and mouse hepatocytes^53^. ESRP2 is a key splicing factor in the liver that maintains the non-proliferative, mature phenotype of adult hepatocytes^14^. Following injury, ESRP2 activity is transiently suppressed to activate a fetal splicing program, which attenuates Hippo signaling and facilitates liver regeneration ^38^. But persistent suppression of ESRP2 in chronic liver disease is detrimental and affects liver functioning and recovery^82, 83^. Therefore, delineating mechanisms regulating ESRP2 expression/activity in developing and diseased livers is critically important. In this study, we provide multiple lines of evidence that ESRP2 activity in maturing hepatocytes is synchronized through overlapping transcriptional, post-transcriptional, and subcellular localization mechanisms.

Whether liver cells exhibit fetal or adult phenotypes is controlled by families of factors that regulate the synthesis, splicing, stability, and translation of mRNAs^14, 84–86^. This biology permits fetal liver cells to be enriched with proteins that allow functions required to accomplish liver organogenesis (e.g., proliferation, migration, morphogenesis) while limiting the expression of proteins that are unnecessary *in utero* due to redundant maternal metabolic, detoxification, and excretory functions. However, after birth, hepatocytes must acquire the proper repertoire of proteins that both enable them to survive *ex utero* and assume mature liver functions that are vital for the life of the organism. For instance, sequential changes in alternative splicing in the postnatal period help generate new proteoforms to support hepatocyte maturity^14, 87^. Consistent with our previous smaller-scale study^14^, ESRP2 deficiency prevented the fetal-to-adult switch in splicing for hundreds of RNA transcripts that normally shift their splicing patterns in the suckling-to-weaning transition period. We also demonstrated that premature ESRP2 activation in the livers of newborn pups forces an earlier-than-normal onset of adult RNA processing program, accelerating hepatocyte maturation and polyploidization.

To identify a mechanistic connection between ESRP2 and how it regulates hepatocyte ploidy, we sought to determine its direct targets. For this, we developed a mouse model using CRISPR-Cas9 mutagenesis wherein the endogenous *Esrp2* allele was FLAG-tagged, enabling a direct pulldown of ESRP2-bound RNAs. eCLIP-seq provided a network of high-confidence, *in vivo* ESRP2-RNA interactions at single-nucleotide resolution. We found that ESRP2 binds extensively to the intronic and 3’-UTR segments of RNA transcripts and prefers a bipartite motif wherein the binding site is UG-rich followed by a Poly G-tract. Integration of eCLIP-seq and RNA-seq datasets from *Esrp2* KO and OE livers deconstructed the ESRP2 splicing-regulatory-network, revealing its direct RNA targets in hepatocytes. Surprisingly, we detected robust ESRP2 binding to the liver-specific *miR-122* host gene upstream of the region that encodes the miRNA stem-loop sequence. Furthermore, we noted that mature miR-122 levels closely mirror ESRP2 protein expression in the postnatal period. ESRP2 deletion caused an accumulation of unprocessed primary miR-122 transcripts, with a corresponding decrease in processed pre-miR-122 and miR-122 transcripts. This was particularly interesting because a postnatal surge in miR-122 is required for the physiological polyploidization of hepatocytes^21, 88^. Additional inquiry revealed that in ESRP2 deficient hepatocytes there is a selective and significant increase in the levels of miR-122 target genes, especially cytokinesis factors. Importantly, *in vivo* and primary cell culture experiments demonstrated a direct dependence of miR-122 and its gene regulatory network on ESRP2. These experiments further indicated that timed activation of ESRP2 boosts the miR-122-driven program of cytokinesis failure and the postnatal expansion of binucleate hepatocytes.

In conclusion, this study provided a deep and mechanistic understanding of the function and regulation of an RNA binding protein in liver maturation. We also uncovered an ESRP2-miR-122 axis that coordinates the postnatal switch from cytokinesis to endo-replication, ensuring proper onset and extent of hepatocyte polyploidization.

### Limitations of the study

We acknowledge that this study has certain limitations and highlight some of them here. When we ectopically expressed ESRP2 in neonatal mice livers and observed premature appearance of specific molecular and cellular signatures, we concluded that ESRP2 is sufficient to induce early maturation of hepatocytes. In retrospect, we realize this is a complex statement. The singular mechanism by which ESRP2 promotes earlier-than-normal hepatic maturation is unclear and assumes that other determinants of hepatocyte maturity are adequately expressed. Also, we found that ESRP2 preferably binds 3’-UTRs over introns; but we do not yet know if or how ESRP2 binding to the 3’-UTRs affects the expression of corresponding transcripts and whether that aids hepatocyte polyploidy and/or maturity.

We showed that ESRP2 binds the primary miR-122 host gene transcript and stimulates its processing to support postnatal binucleation/polyploidization of hepatocytes. While we demonstrated that the increased incidence of binucleation in ESRP2 expressing primary hepatocytes is dependent on miR-122, we cannot rule out the possibility that other intermediary factor(s) are also involved. For example, loss of ESRP2 does not completely inhibit miR-122 production or entirely block hepatocyte polyploidization. Finally, it is plausible that as a maturity factor, ESRP2 supports the general maturation of the hepatic environment, making it more permissive to miR-122 expression without the need for a direct interaction. Future studies involving deletion of ESRP2 binding sites within the primary miR-122 host transcript *in vivo* could help clarify this alternative possibility.

## KEY RESOURCES TABLE

| REAGENT or RESOURCE | SOURCE | IDENTIFIER |
| --- | --- | --- |
| **Antibodies** |  |  |
| ESRP2, mouse, Primary | Rockland | 210-301-C31 |
| TBP, mouse, Primary | Thermo Fisher | MA5-14739 |
| GAPDH, rabbit, Primary | Sigma-Aldrich | G9545-200UL |
| HNF4A, mouse, Primary | Abcam | ab41898 |
| HNF4A, rabbit, Primary | Cell signaling Tech | 3113 |
| FLAG M2, mouse, primary | Sigma Aldrich | F1804 |
| Beta-Catenin, rabbit, primary | Cell signaling Tech | 9562 |
| Goat Anti-Mouse IgG, Secondary HRP | Bio-Rad | 1721011 |
| Goat Anti-Rabbit IgG, Secondary HRP | Thermo Fisher | 31460 |
| Goat Anti-Mouse IgG, Secondary 488 | Thermo Fisher | 35503 |
| Goat Anti-Mouse IgG, Secondary 594 | Thermo Fisher | 35511 |
| Goat Anti-Rabbit IgG, Secondary 488 | Thermo Fisher | 35553 |
| Goat Anti-Rabbit IgG, Secondary 594 | Thermo Fisher | 35561 |
| **Bacterial and Virus Strains** |  |  |
| Stellar™ Competent Cells | Takara Bio | 636766 |
| **Deposited Data** |  |  |
| RNA-Sequencing | This study | GEO: GSE217391 and GSE151309 |
| **Experimental Models: Organisms/Strains** |  |  |
| Germline *Esrp2*<sup>KO/KO</sup> | PMID: 26371508 |  |
| TRE-ESRP2<sup>Tg/wt</sup> | PMID: 30250226 |  |
| ApoE-rtTA<sup>Tg/Tg</sup> | PMID: 17889654 |  |
| 2xFLAG-ESRP2<sup>Tg/wt</sup> | This study |  |
| ESRP2<sup>flox/flox</sup> | This study |  |
| **Software and Algorithms** |  |  |
| Trimmomatic (version 0.38) | PMID: 24695404 | RRID:SCR_011848 |
| STAR (version 2.4.2a) | PMID: 23104886 | RRID:SCR_015899 |
| Cuffdiff (version 2.2.1) | PMID: 22383036 | RRID:SCR_001647 |
| rMATS (version 3.2.5) | PMID: 25480548 | N/A |
| DAVID (version 6.8) | PMID: 12734009 | RRID:SCR_001881 |
| HOMER (version 3.12) | PMID: 20513432 | RRID:SCR_010881 |
| MEME (version 5.0.4) | PMID: 20513432 | RRID:SCR_001783 |
| CTK (version 1.0.8) | PMID: 27797762 | N/A |
| SCENIC version 1.3.1 | PMID: 28991892 | N/A |
| Seurat version 5.0.3 | PMID: 37231261 | N/A |
| Monocle version 2.26.0 | PMID: 28114287 | N/A |

## STAR*METHODS

### LEAD CONTACT AND MATERIALS AVAILABILITY

Further information and requests for resources and reagents should be directed to and will be fulfilled by the Lead Contact, Auinash Kalsotra.

### EXPERIMENTAL MODEL AND SUBJECT DETAILS

#### Experimental Animals

National Institutes of Health (NIH) and institutional guidelines were followed in the use and care of laboratory animals and experimental protocols were performed as approved by the Institutional Animal Care and Use Committee (IACUC) at UIUC. Mice were housed on a standard 12-hour-light/dark cycle (18-23 °C ambient temperature; 40-60% humidity) and were allowed *ad libitum* access to water and a normal chow diet (2918 Envigo Teklad). Genomic DNA was derived from tail biopsies and genotyping was performed using standard procedures. Both male and female mice were used in this study. Whole liver tissues and hepatocytes were isolated from mice following guidelines for euthanization and/or anesthesia.

Generation of germline *Esrp2* KO^14^ and Dox-inducible, hepatocyte-specific ESRP2 overexpression (TRE-ESRP2; ApoE-rtTA)^38^ mice were described previously. All mice reported in this study were F1 progeny of TRE-ESRP2 and ApoE-rtTA transgenic mice mating, and always hemizygous for both transgenes and the ApoE-rtTA transgene. For conditional knockout of *Esrp2*, flox/flox (f/f) animals were generated wherein exons 3-13 of the *Esrp2* gene locus were floxed in mice. The targeting vector for *Esrp2* was designed in collaboration with the Penn Gene Targeting Core. The vector featured a loxP site in intron 2 upstream of exon 3, a neomycin selection cassette flanked by FRT sites in intron 14 followed by another loxP site. The targeting vector was electroporated into V6.5, hybrid C57BL6/129Sv mouse ES cells and neomycin-resistant clones were isolated, screened by Southern blotting, injected into Balb/c blastocysts, and implanted in pseudo-pregnant females. Chimeras were generated and males were crossed to C57BL/6J females. Germline transmission was confirmed by PCR/Southern blots and the conditional allele was created by crossing the chimeric male with a B6(C3)-Tg(Pgk1-FLPo)10Sykr/J female to remove the neomycin cassette. Progeny were screened for removal of the neomycin cassette by PCR analysis. Heterozygous *Esrp2* ^f/wt^ mice were crossed to produce homozygous *Esrp2* ^f/f^ conditional mice, with no apparent phenotype, and to maintain the line. For acute knockout studies, *Esrp2* ^f/f^ mice were injected via tail vein with either AAV8-TBG-CRE or AAV8-TBG-GFP virus (Vector Bio-labs Inc., viral titer of 5 × 10^11^ genome copies) wherein the CRE or GFP proteins were expressed only in hepatocytes **(Figure S2B)** due to the use of TBG promoter (Thyroxine binding globulin, Hepatocyte-specific expression).

For generating N-terminal 2xFLAG tagged ESRP2 transgenic mice at UIUC transgenic mouse core facility, crRNA (gRNA sequence - CCGCGTGTGTGGCCGAGCGCCC), tracrRNA, espCas9 and an oligonucleotide repair template encoding two tandem copies of FLAG epitope tag (5’-GATTACAAGGATGACGACGATAAGGCAGATTACAAGGATGACGACGATAAG-3’) were injected into C57BL/6J embryos to derive F0 progeny. To identify the positive founder animals, mice were genotyped using standard PCR on tail-clip-derived genomic DNA. The germline incorporation of the 2xFLAG tag was further confirmed by Sanger sequencing of the F1 progeny. For eCLIP experiments, heterozygous FLAG ESRP2 ^Tg/wt^ mice were used.

#### Murine Diet Scheme

For all experiments, the nursing female was fed a standard diet containing 6.0 g/kg Doxycycline (Teklad, Envigo) starting on the day the pups were born. This diet was continued till postnatal day 21, followed by a 0.1 g/kg Doxycycline diet till the end of the experiment.

### METHOD DETAILS

#### Hepatocyte Isolation

Hepatocytes from mouse livers of indicated genotypes were isolated using adapted protocols from previously described methods^14, 38^. Briefly, P14 or Adult (P56) mice were anesthetized in a chamber supplied with Isoflurane and Oxygen (2.5% Isoflurane in oxygen, 1L/min), and maintained on the anesthetic during the procedure. The liver was perfused via cannulation of the portal vein with 50 mL of Solution A (1x HBSS (w/o Ca^2+^ and Mg^2+^), 0.5 mM EDTA) followed by 50 mL of Solution B (1x HBSS (w/ Ca^2+^), 5.4 mM CaCl_2_, 0.04 mg/mL Soybean Trypsin Inhibitor, and 3000 Units of Collagenase type IV (Worthington Chemicals)). Subsequently, the liver was massaged in a Petri dish containing 1x HBSS to release cells from the liver capsule, which were passed successively through and 70 and 40 μm filters to obtain a single-cell suspension. The cells were then centrifuged at 50 x g for 5 min (4°C) to separate live hepatocytes from non-parenchymal cells (NPCs) and dead cells. Hepatocytes were further washed 3 times in 1x HBSS, flash frozen in liquid N_2_, and stored at −80°C till further use.

#### Bulk RNA-seq and Data Analysis

Hepatocyte RNAs were isolated using RNeasy tissue mini-kit (Qiagen) and the RNA quality was assessed with Agilent Bioanalyzer followed by quantification by Qubit Fluorimeter at the Functional Genomics Core in Roy J. Carver Biotechnology Center, UIUC. Hi-Seq libraries were prepared, and 150 bp paired-end Illumina sequencing was performed on a HiSeq 4000 or NovaSeq 6000 at the High Throughput Sequencing and Genotyping Unit, UIUC. RNA-seq reads were processed for quality/read length using Trimmomatic (version 0.38) and then aligned to the mouse genome (mm10) using STAR (version 2.4.2a). RNA abundance levels were determined as Transcript per million (TPM) using count and differential expression values obtained from DESeq2 (version 1.8.2), and HTseq (version 0.6.1). RNAs were considered differentially expressed following imposed cutoff clearance (False Discovery rate (FDR) < 0.05, |Log_2_ (Fold Change)| > 1). Differential splicing analysis was carried out using rMATS (version 3.2.5), and significant events were identified with imposed cutoffs (FDR < 0.10, Junction read counts ≥ 10, percent spliced In ≥ 15%). Motif analysis for differentially spliced exons was performed using rMAPS with default parameters, and putative motifs were identified as described previously ^38^. Gene ontology analysis was performed using DAVID (version 6.8) and mapped with “Enrichment Maps” plugin in Cytoscape. All expressed genes with Transcript per million (TPM) >1 served as background, and the biological functions were analyzed with three pathways (Biocarta, Kegg, and Panther). Functional clustering was performed, and the top clusters (p < 0.05) represented. For Exon Ontology analysis, regulated exons in maturing hepatocytes were identified, and their mouse coordinates (mm10) were converted to corresponding human exons in the hg19 annotation using UCSCliftover with minimum ratio of bases matching as 0.8. The selected exons in hg19 were further checked for gene identity match to the mouse exon’s parent gene. Once verified, the exon sets were analyzed for ontological processes using Exon ontology, and FasterDB packages^89^.

#### Isolation of Liver Cells, Single-cell Library Preparation and Sequencing

After the mouse was anesthetized through continuous inhalation of 2% isoflurane, a 5-cm long incision was made in the abdomen to expose the portal vein and inferior vena cava. To perfuse the liver, the portal vein was cannulated, and ∼30 ml of solution I (1x HBBS without Ca2+/Mg2+ with 1 mM EDTA, 37 °C) was passed followed by ∼40-50 ml of solution II (HBSS with 0.54 uM CaCl2, 40 ug/ml Trypsin Inhibitor, 15 mM HEPES PH 7.4 and 6000 units of collagenase IV, 37 °C). The liver was excised and massaged in washing solution (DMEM + Ham’s F12 (50:50) with 5% FBS and 100 U/ml Penicillin/Streptomycin, 4 °C) to release cells from the liver capsule. The cell suspension was then passed through 40 um filters to remove doublets/undigested tissue chunks, pelleted by centrifugation at 350 x g for 4 min at 4 °C to remove debris and resuspended in 15 ml wash buffer. Cells were counted with an automated hemocytometer, and ∼1-1.5 million cells were processed for library preparation. MACS Dead Cell Removal Kit (Miltenyl Biotec) was used to remove dead cells and obtain single-cell suspensions with high viability. Following this, single-cell sequencing libraries were prepared (from a pool of two mice livers for each time point) using the 10X Genomics Chromium Single Cell 3’ Kit v3 and sequenced with Illumina NovaSeq 6000 on a SP/S4 flow cell to obtain 150 bp paired reads.

#### Single-cell RNA-seq Analysis

Single-cell libraries produced over one and a half billion reads. We used Cell Ranger v3.1 pipelines from 10X Genomics to align reads and produce feature barcode matrices. Seurat v4.0^90^ was used for QC and analysis of individual feature barcode matrices were further integrated after removing batch-specific effects using Harmony v1.2.0^91^. Data was log-normalized, scaled, and clustered after PCA analysis. Hepatocyte and NPC clusters were identified based on the expression of multiple marker genes. To identify cell types within the subset of NPCs, they were further subjected to unsupervised UMAP clustering.

Monocle v2.0 was used to perform pseudotime analysis, according to the online documentation^92^. The CellDataSet class monocle objects were made from log normalized Seurat object containing cells under consideration. Dimensionality reduction was performed with the DDRTree algorithm, and the 1500 most significant deferentially expressed genes were used to perform pseudotime ordering and obtain cell trajectories. Genes with expression patterns co-varying with pseudotime were determined by the ‘differentialGeneTest()’ module, clustered and plotted using the ‘plot_pseudotime_ heatmap()’ module. Expression patterns in clusters were distinguished as upregulated/downregulated along the pseudotime, and gene ontology analysis was performed using DAVID 6.8^93^ to identify biological pathways that were up or downregulated along the pseudotime.

SCENIC^45^ pipeline was used to estimate the AUCell Score activity matrix from the log-normalized Seurat object containing the subset of hepatocytes. Unlike the standard SCENIC workflow where this AUCell score activity matrix is binarized by thresholding to generate binary regulon-activity matrix, we retained the full AUCell score for all further analysis. UMAP plots, heatmaps and violin plots demonstrating regulon activities based on AUCell scores were made in Seurat 3.1. AUCell scores were plotted over pseudotime cell-trajectories using Monocle 2.0.

#### Polysome Gradient Fractionation

For isolation of polysomes from P14 and P28 livers, a modified perfusion protocol was developed wherein the animals were first perfused via portal vein cannulation with 75mL of Solution I (300 ug/mL Cycloheximide in 1x HBSS (w/ phenol red)) at rate of 3.5 mL/min. EDTA was excluded from Solution I due to its dissociative effects on ribosomes and associated complexes. Instead, liver perfusion and hepatocyte wash buffers were supplemented with Cycloheximide (300 ug/mL) to arrest translating ribosomes. For E18 and P7 timepoints, hepatocytes were isolated with a digestion protocol wherein all solutions contained cycloheximide as reported previously^38, 39^, since perfusions at these timepoints are not feasible.

Purified hepatocytes were thawed in polysome lysis buffer (10 mM Tris-HCl at pH 8.0, 150 mM NaCl, 5 mM MgCl2, 1 mg/mL heparin, 1% Nonidet-P40, 0.5% deoxycholate, 40 mM dithiothreitol, 1 U/mL SUPERaseIn RNase inhibitor [Thermo Fisher], and 150 µg/mL cycloheximide) and lysed through gentle pipetting. The cell nuclei and cell debris were removed by centrifugation (12,000 *g*, 1 min, at 4°C). The supernatant was transferred to a fresh tube and then centrifuged again to remove remaining cell debris and organelles (16,000*g*, 7.5 min, at 4°C). The resulting supernatant was transferred to a fresh tube where 400 µL was layered onto a 10-mL linear sucrose gradient (15%–45% sucrose [w/v] made using a Biocomp Gradient Master) and centrifuged in a SW41Ti rotor (Beckman) for 125 min at 38,000 rpm and 4°C, as previously described. Polysome profiles were recorded using a UA-6 absorbance (ISCO) detector at 254 nm and fractions were collected along the gradient corresponding to the RNP, monosome, light polysomes (2-4 ribosomes), and heavy polysomes (5+ ribosomes). RNAs for downstream sequencing and validations were purified using an adapted protocol^94^. The sucrose gradient fractions were mixed with 0.05 volume of 3M Sodium Acetate plus 2 μL Glycogen followed by addition of 2 volumes of 100% EtOH and stored at −80 C overnight to precipitate RNA/protein. The pellets were suspended in molecular grade water, and after DNase digestion, samples were phase separated using 3 volumes of Acid phenol: Chloroform (5:1) mixture. The aqueous phase was phase separated again with Chloroform: Isoamyl Alcohol (24:1). After that the aqueous phase was mixed with 0.1 volume 3M Sodium Acetate and 2 uL glycogen, and precipitated with 2.5 volumes EtOH overnight at −80 C. The pelleted RNAs were then used for polyA+ selected deep sequencing or qRT-PCR.

#### Histology and Immunofluorescence Staining

Liver tissues were fixed in 10% buffered Formalin for 24 hours, processed, and embedded in paraffin. 5 μm tissue sections were cut, and deparaffinized in Xylene and moved through serial alcohol washes (100, 95, 80, 50%) to rehydrate. For H&E staining, sections were washed in Hematoxylin (2 min) and Eosin (1 min) successively, cover slipped with permount medium, and imaged on Hamamatsu Nanozoomer at the IGB core facility, UIUC. For Immunofluorescent staining, deparaffinized and rehydrated sections were antigen retrieved in a slow cooker at 120°C for 10 minutes. The retrieval was performed in either TE (Tris-EDTA pH 8.0) or citrate buffer (10 mM Sodium citrate, 0.05% Tween 20, pH 6.0). The sections were then washed in buffer A (TBS + 0.05% Triton X-100) and blocked (2 h) in 10% Normal Goat Serum (NGS) containing 1% BSA at room temperature (RT). Primary antibodies were applied to the sections at standardized concentrations and incubated overnight at 4°C. Next, the sections were washed in buffer A, and secondary fluorescent antibodies applied for 1 hour at RT. Lastly, ToPro Nuclear stain was applied for 15 minutes at RT and the sections were cover slipped using CC aqueous mounting media. All sections were imaged on a Zeiss LSM 710 microscope at the IGB core facility, UIUC. To stain for nascent DNA synthesis in maturing livers, mice were treated with a 2-hour pulse of EdU (5-ethynyl-2’-deoxyuridine) administered intra-peritoneally (50 μg/g of body weight). Mice were sacrificed, and the livers were harvested, and paraffin embedded. 5 μm sections were deparaffinized, rehydrated, and antigen retrieved in citrate buffer (10 mM Sodium citrate, 0.05% Tween 20, pH 6.0). The sections were then washed in buffer A (TBS + 0.05% Triton X-100) and blocked (2 h) in 10% NGS and 1% BSA at RT. EdU labeled DNA was stained with Alexa Fluor 488 using Click-iT EdU Alexa Fluor kit (Thermo Fisher) as per manufacturer’s protocol.

#### Protein Isolation and Western Blot Analysis

Protein isolation was performed on frozen liver tissue or isolated hepatocyte fractions using homogenization in bullet blender followed by sonication. The homogenization buffer contained 10 mM HEPES-KOH, pH 7.5, 0.32 M sucrose, 5 μM MG132, 5 mM EDTA, and Pierce proteinase inhibitor tablet (1 tablet/ 20 mL buffer volume). Prior to sonication, 20% Sodium Dodecyl sulfate (SDS) to a final concentration of 1% (v/v) was added. Protein concentration was measured using Thermo scientific BCA assay kit. Approximately 50 μg of total protein sample was loaded onto a 10% SDS-PAGE gel, and after separation by gel electrophoresis, proteins were transferred onto a PVDF membrane. Membranes were visualized for equal loading using Ponceau (PonS) staining solution (0.5% w/v PonS, 1% Acetic acid), and then blocked using 5% milk powder (w/v) in TBST (Tris-buffered saline, 0.1% Tween 20) for 2 hours at RT. Blots were incubated in primary antibodies at respective concentrations overnight at 4°C. After TBST washes, blots were incubated in HRP-conjugated secondary antibodies for 1 hour at RT and developed using Clarity Western ECL kit (BioRad).

#### Gene Expression and Splice Isoform Analysis

Total RNAs from primary hepatocytes, or mouse livers were isolated using either RNeasy kit (Qiagen) or TRIzol reagent (Life Tech). Following DNase I treatment, 5 μg of RNA was reverse transcribed to cDNA using random hexamers and Maxima Reverse transcriptase kit (Thermo Scientific). The cDNA was diluted to a final concentration of 25 ng/μL and used for quantitative real time PCR (qRT-PCR) or splicing-sensitive RT-PCR assays. Downstream analyses for PSI and log2 fold change calculations were performed as described previously^14^. For quantifying expression of U6 snRNA and miR-122, Taqman assays 001973 and 002245 (Life Technologies) were used respectively.

#### Enhanced Crosslinking and Immunoprecipitation (eCLIP)

Isolated hepatocytes from FLAG ESRP2 ^Tg/wt^ mice were suspended in 1xPBS and crosslinked with 1 pulse of 400 mJ/cm^2^ of 254 nm UV radiation to stabilize RBP–RNA interactions. Subsequent immunoprecipitation (IP) of FLAG ESRP2-RNA complexes, RNA isolation, library preparation and sequencing were performed as described previously^46^. Briefly, crosslinked cells were lysed in buffer and sonicated, followed by treatment with RNase I (Thermo Fisher) to fragment RNA. FLAG M2 antibody (Sigma Aldrich, F1804) was pre-coupled to anti-mouse IgG Dynabeads (Thermo Fisher, 11201D), added to the lysates, and incubated 3 hours at 4 °C. Prior to IP washes, 2% of sample was removed to serve as the paired input sample. For IP samples, high- and low-salt washes were performed, after which RNA was dephosphorylated with FastAP (Thermo Fisher) and T4 PNK (NEB) at low pH, and a 3′ RNA adaptor was ligated with T4 RNA ligase (NEB). 15% of IP and input samples were run on an analytical 4-12% PAGE gel, transferred to PVDF membrane, blocked in 5% dry milk in TBST, incubated with ESRP2 antibody (Rockland, 210-301-C32), washed, incubated with HRP-conjugated anti-mouse secondary (BioRad, 1706516), and visualized with chemiluminescence imaging to validate successful IP. The remaining IP and input samples were run on an 4-12% PAGE gel and transferred to nitrocellulose membranes, after which the region between 78-155 kDa was excised from the membrane, treated with proteinase K (NEB) to release RNA, and concentrated by column purification (Zymo). Input samples were then dephosphorylated with FastAP (Thermo Fisher) and T4 PNK (NEB) at low pH, and a 3′ RNA adaptor was ligated with T4 RNA ligase (NEB) to synchronize with IP samples. Next, reverse transcription was performed with AffinityScript (Agilent), followed by ExoSAP-IT (Affymetrix) treatment to remove unincorporated primer. RNA was then degraded by alkaline hydrolysis, and a 3′ DNA adaptor ligated with T4 RNA ligase (NEB). qRT-PCR was used to determine the required amplification cycles, followed by PCR with Q5 (NEB) and gel electrophoresis to size-select the final library. Libraries were sequenced on the NovaSeq6000 platform (Illumina). eCLIP was performed on IP from two independent samples, along with paired size-matched input before the IP washes.

#### Primary Hepatocyte Cultures

Primary hepatocyte culture experiments were performed as described previously^21^. Briefly, hepatocytes from wildtype P14-15 C57BL/6 mice were seeded at 40-50k density/well in 24-well Primaria Cell Culture plates (Corning) in growth media: DMEM-F12 with 15 mM HEPES (Corning), 5% FBS (Atlanta Biologicals), Antibiotic-Antimycotic Solution (Corning) and ITS Supplement (containing 5 μg/ml insulin, 5 μg/ml transferrin and 5 ng/ml sodium selenite; Roche). After 16h, growth media was replaced with fresh media lacking Antibiotic-Antimycotic Solution, and cells were transfected with 25 nM miRIDIAN miR-122 mimic (Fisher, C-300591-05-0005), miRIDIAN negative control (Fisher, CN-001000-01-05) (Life Technologies) or 10 nM mirVana miR-122 inhibitor (Cat no. 4464084, Assay ID MH11012, Thermo Fisher) using Lipofectamine 2000. Six hours after transfection, media was replaced with fresh growth media containing 0.5% FBS. Morphological changes were documented in cultures fixed with 4% PFA. The numbers of mono- and bi-nucleate hepatocytes were quantified on day 4.

#### Ploidy Analysis

For detection of ploidy populations, freshly isolated primary hepatocytes were washed twice in PBS, adjusted to a density of 1-2 million cells/ml and incubated on ice with 2 μl/ml Fixable Viability Dye (FVD) eFluor 780 (eBioscience, San Diego, CA). Following 2 washes with PBS, hepatocytes were fixed with 2% PFA, washed an additional 2 times with PBS, and incubated with permeabilization buffer (0.1% saponin and 0.5% BSA in PBS) + 15 μg/ml Hoechst 33342 (Life Technologies, Carlsbad, CA). Cells were washed twice and stored in PBS until flow cytometry analysis. We also detected Ki-67 in hepatic ploidy populations. Hepatocytes were prepared using the same methods as above except following FVD 780 incubation they were fixed with Foxp3 fixation buffer (eBioscience) at room temperature, washed twice with permeabilization buffer (eBioscience) and incubated in permeabilization buffer with mouse anti-Ki-67 eFluor 660 antibody (Invitrogen, Carlsbad, CA) + 15 μg/ml Hoechst 33342. Cells were washed twice with PBS and stored in permeabilization buffer until flow cytometry analysis. Cells were analyzed with an LSR II flow cytometer (BD Biosciences, Franklin Lakes, NJ) running BD FACSDiva™ SoftwarepH v9.0. FACS plots were generated using FlowJo 10.8.2 (FlowJo LLC, Ashland, OR).

#### Statistical Analysis

All quantitative experiments (qRT-PCR, western blots, immunofluorescence) have at least three independent biological repeats. The results were expressed with mean and standard deviation, unless mentioned otherwise. Differences between groups were examined for statistical significance using unpaired t-test with Welch’s correction (for two groups), or one/two-way ANOVA for more than two groups using the GraphPad Prism 6 Software. All p-values are listed on respective comparisons in plots; p-value <0.05 was deemed significant.

#### Accession numbers

All raw RNA-seq and eCLIP-seq data files are available for download from NCBI Gene Expression Omnibus (http://www.ncbi.nlm.nih.gov/geo/) under accession numbers GSE217391 and GSE151309.

## SUPPLEMENTAL INFORMATION

Not applicable.

## ACKNOWLEDGEMENTS

We thank the members of the Kalsotra laboratory for valuable discussions and comments on the manuscript. This work was supported through the NIH grants R01AA010154, R01HL126845, R21HD104039 (to A.K.), R01DK103645 (to A.W.D.) Chan-Zuckerberg Biohub Chicago Award and Muscular Dystrophy Association Research Grant (to A.K.); the Commonwealth of Pennsylvania (to A.W.D); the NIH Tissue microenvironment training program T32-EB019944 and UIUC Scott Dissertation Completion Fellowship (to S.B.); the Herbert E. Carter fellowship in Biochemistry (to U.V.C.); the NIH pre-doctoral NRSA fellowship F30DK108567 (to W.A.). Transgenic Mouse Core, High-Throughput Sequencing/Genotyping Core, and Histology/Microscopy Core facilities at UIUC and the Pittsburgh Liver Research Center Advanced Cell Tissue and Imaging Core (P30DK120531) supported this project.

## AUTHOR CONTRIBUTIONS

S.B., and A.K. conceived the project and designed the experiments. S.B., J.C., N.B., U.V.C., D.D., J.M.D., S.C., W.A., F.A., and A.W.D. performed experiments. A.W.D., N.G. and R.P.C. provided reagents. S.B. and N.B. performed the bioinformatics analysis. S.B., and A.K. analyzed data, interpreted results and wrote the manuscript. All authors discussed the results and edited the manuscript.

## DECLARATION OF INTERESTS

The authors declare no competing financial interests.

## Notes

### Competing Interest Statement

The authors have declared no competing interest.

