## Supplementary Figures for "ESRP2-microRNA-122 axis directs the postnatal onset of liver polyploidization and maturation"

**Running title:** Posttranscriptional control of liver polyploidy and maturation

**Keywords:** RNA processing, polyploidy, genome editing, Protein-RNA interactions, ESRP2, microRNA, single-cell transcriptomics, eCLIP, hepatocyte maturation

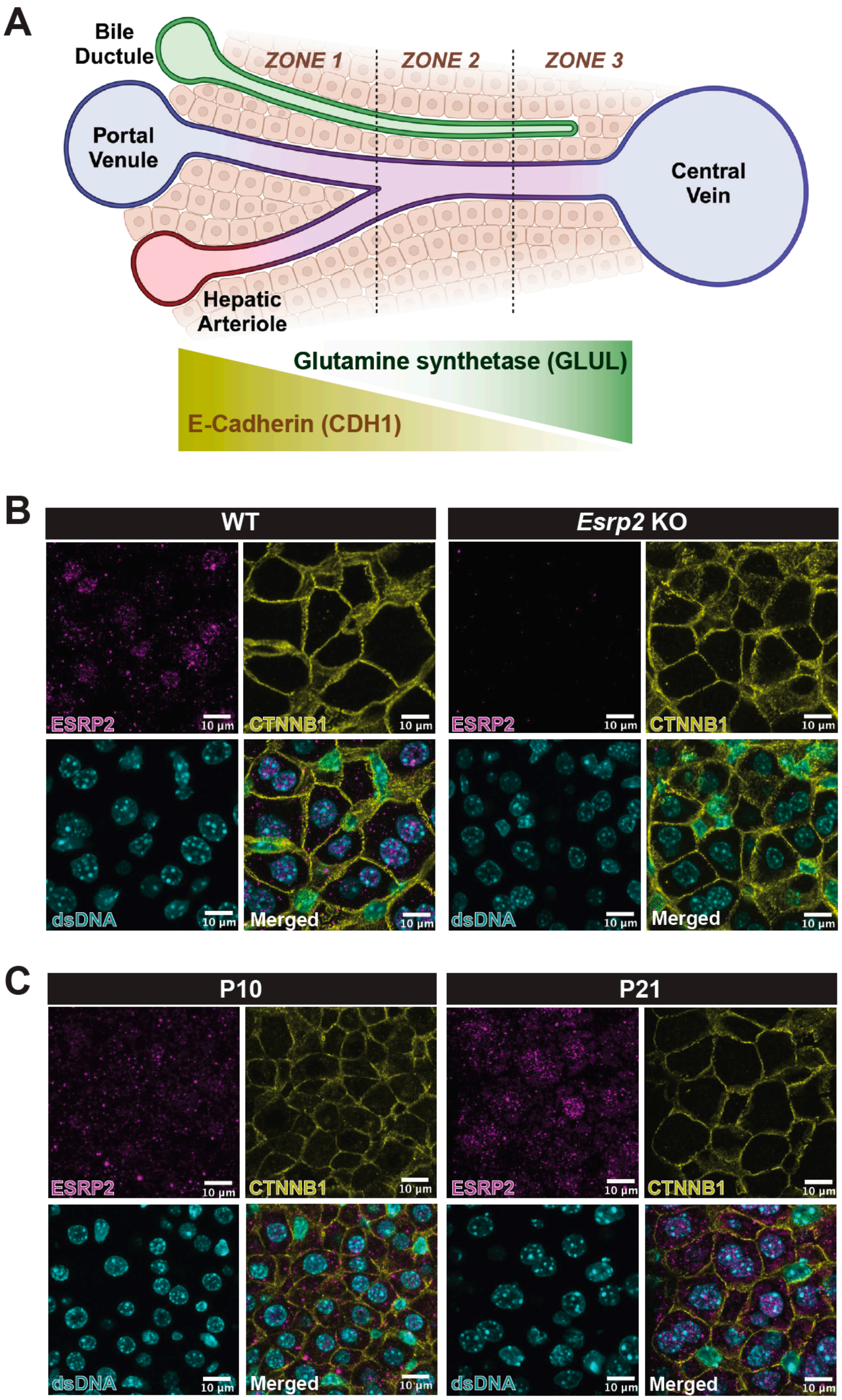


**Supplementary Figure 1. (A)** Schematic for cellular organization within the hepatic lobule, with hepatocytes arranged between the Portal and central veins. E-Cadherin (CDH1) and Glutamine synthetase (GLUL) as zonal markers of hepatocytes surrounding the portal and central vein, respectively. **(B)** Immunofluorescence (IF) staining for ESRP2, and a membrane marker (CTNNB1) in adult wildtype and ESRP2 KO mice livers. **(C)** IF staining for ESRP2, and membrane-specific (CTNNB1) marker in wildtype P10 (LEFT) and P21 (RIGHT) livers. dsDNA was labeled with Hoechst 33342 in (B) and (C).

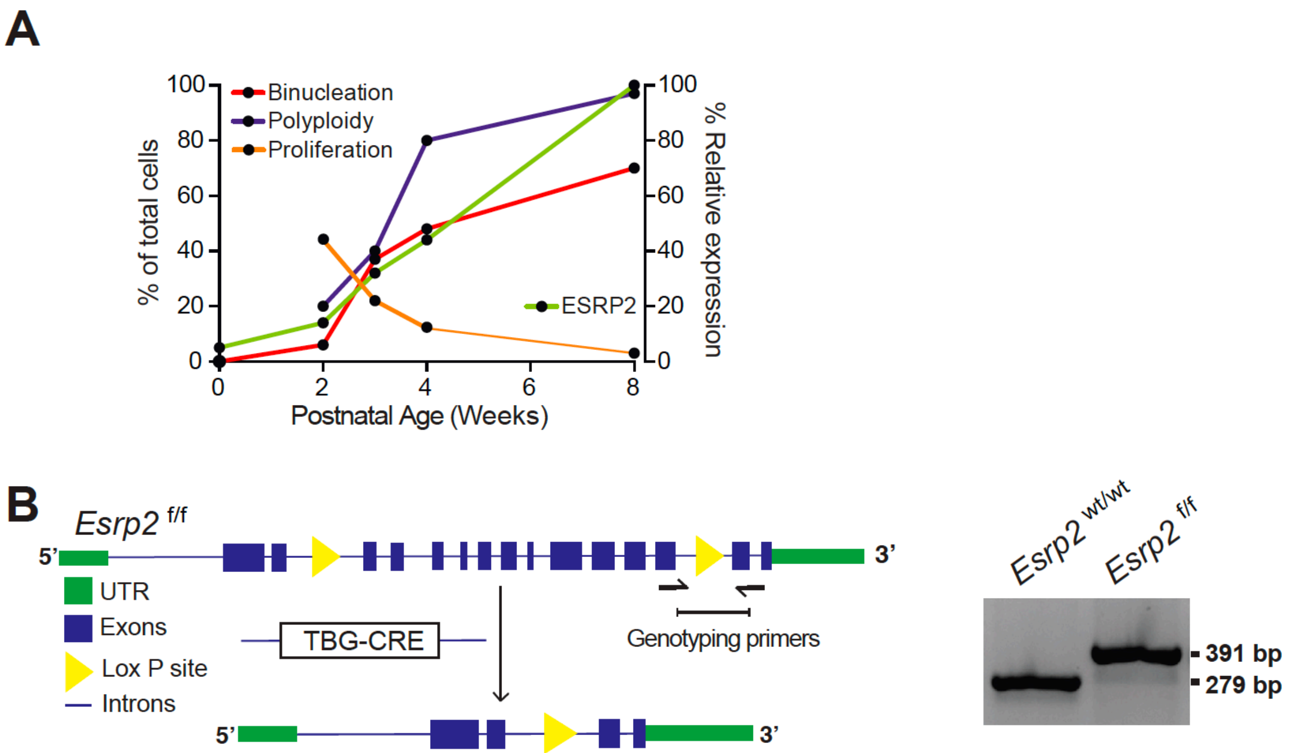


**Supplementary Figure 2. (A)** Relative changes in hepatocyte characteristics (i.e. binucleation, polyploidy and proliferation) and ESRP2 protein levels during postnatal liver maturation in mice. **(B)** (Left) Schematic for conditional deletion allele of *Esrp2* wherein exons 3-13 are surrounded by two Lox P sites. The two arrows indicate genotyping primer positions to identify the downstream Lox P site. (Right) Genotyping gel indicating the presence of the 112 nt insertion harboring the Lox P site between exons 13 and 14 in *Esrp2* $^{f/f}$ mice.

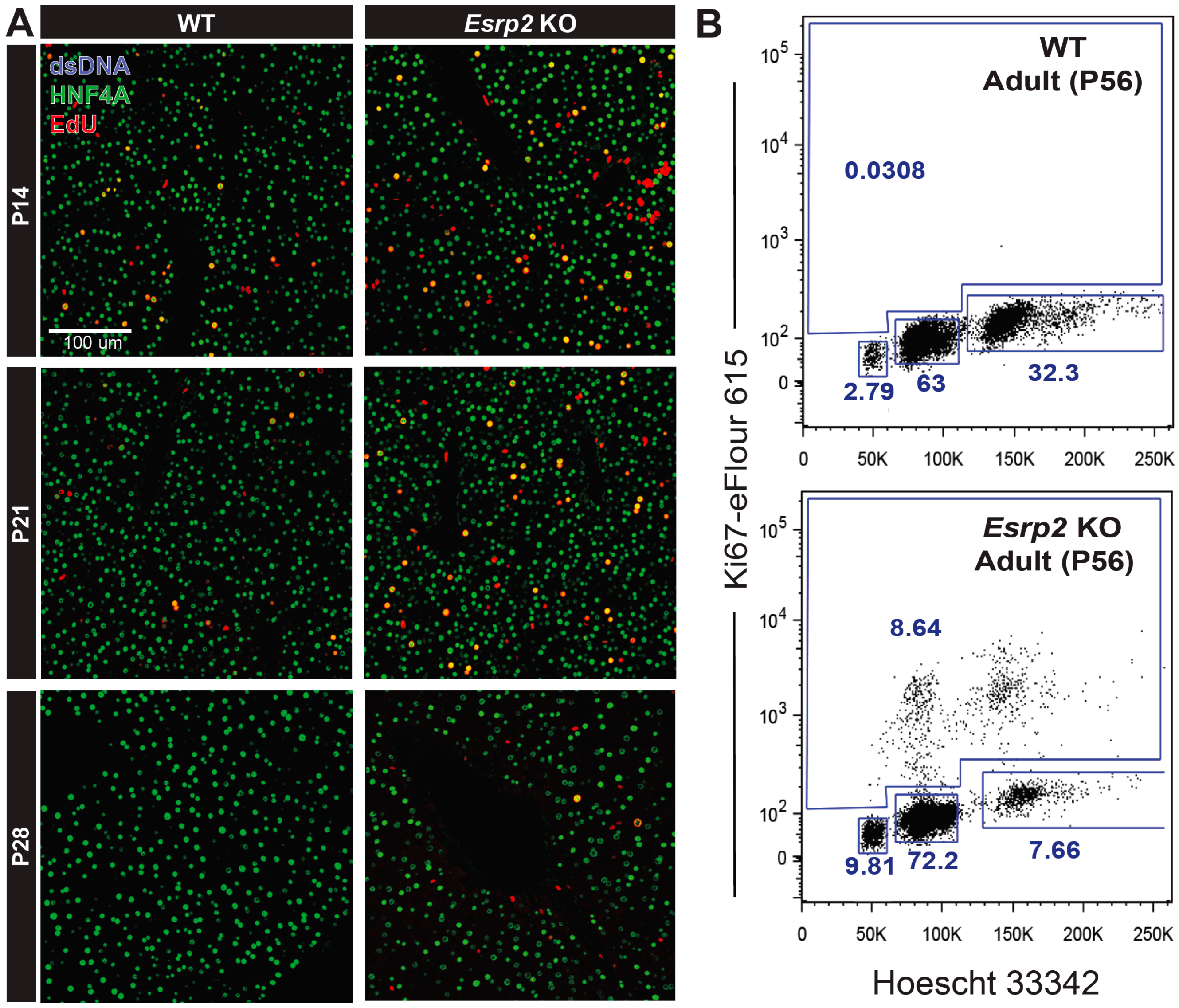


**Supplementary Figure 3. (A)** Representative immunofluorescence images of livers from wildtype (WT), and *Esrp2* knockout (KO) mice at 2, 3 and 4 weeks labeled for EdU incorporation along with HNF4A (Hepatocyte marker) and DAPI (dsDNA marker). **(B)** Flow cytometry derived scatter plots for wildtype (WT), and *Esrp2* knockout (KO) hepatocytes stained for Hoechst 33342 (DNA content marker) and Ki67 (proliferation marker).

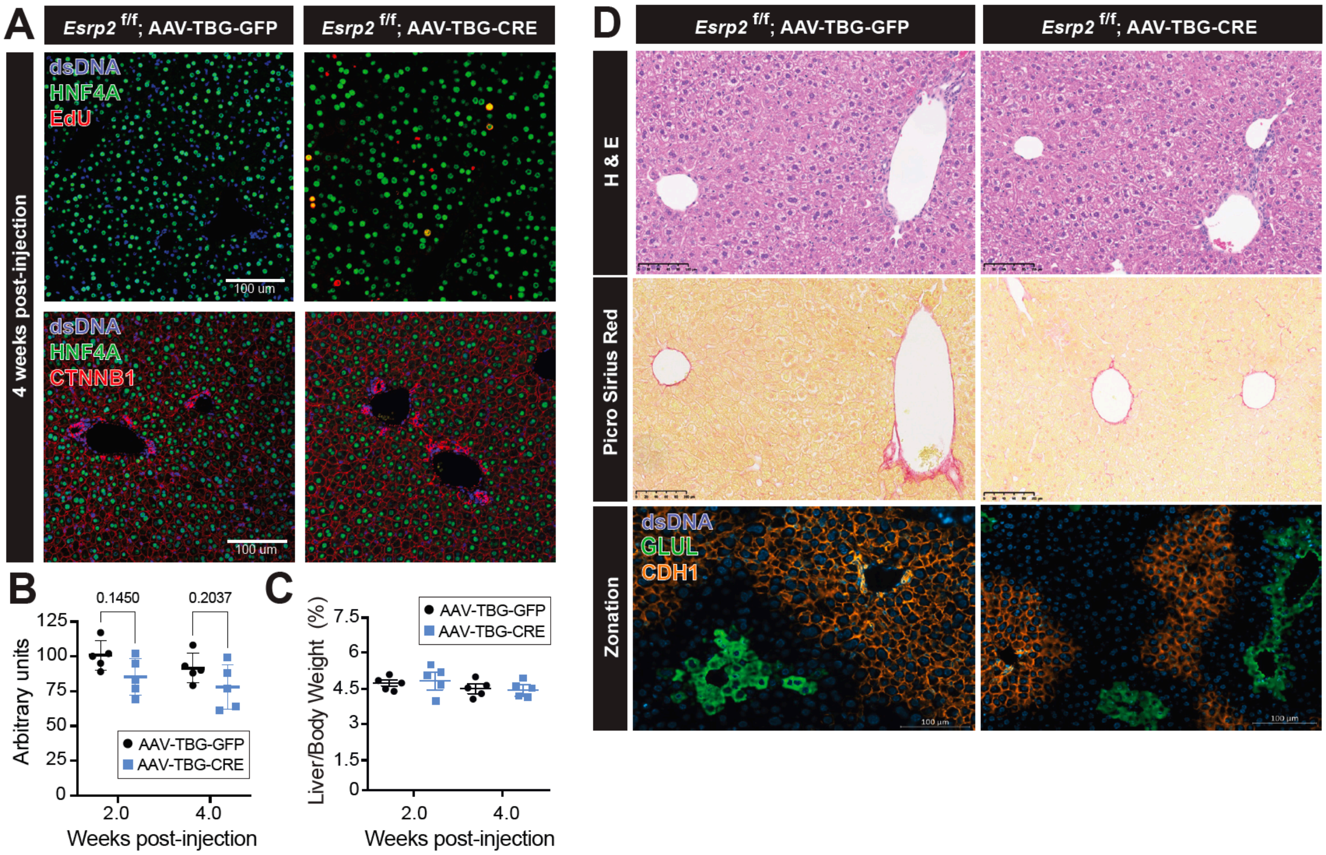


**Supplementary Figure 4. (A)** Representative immunofluorescence images of *Esrp2* $^{f/f}$ livers injected with AAV-TBG-GFP or AAV-TBG-CRE, 4 weeks post-injection stained for (TOP) EdU incorporation, and (BOTTOM) CTNNB1 along with HNF4A (Hepatocyte marker) plus DAPI (nuclear marker). **(B)** Quantification of hepatocyte size from *Esrp2* $^{f/f}$ livers injected with AAV-TBG-GFP or AAV-TBG-CRE at 2- and 4-weeks post-injection. **(C)** Liver-to-body weight quantifications of *Esrp2* $^{f/f}$ livers injected with AAV-TBG-GFP or AAV-TBG-CRE at 2 weeks post-injection. **(D)** Representative histology and immunofluorescence images of *Esrp2* $^{f/f}$ livers injected with AAV-TBG-GFP or AAV-TBG-CRE at 4 weeks post-injection stained for (TOP) Hemotoxylin and Eosin, (MIDDLE) Picro Sirius red, and (BOTTOM) CDH1 and GLUL (zonation markers) and DAPI (nuclear marker).

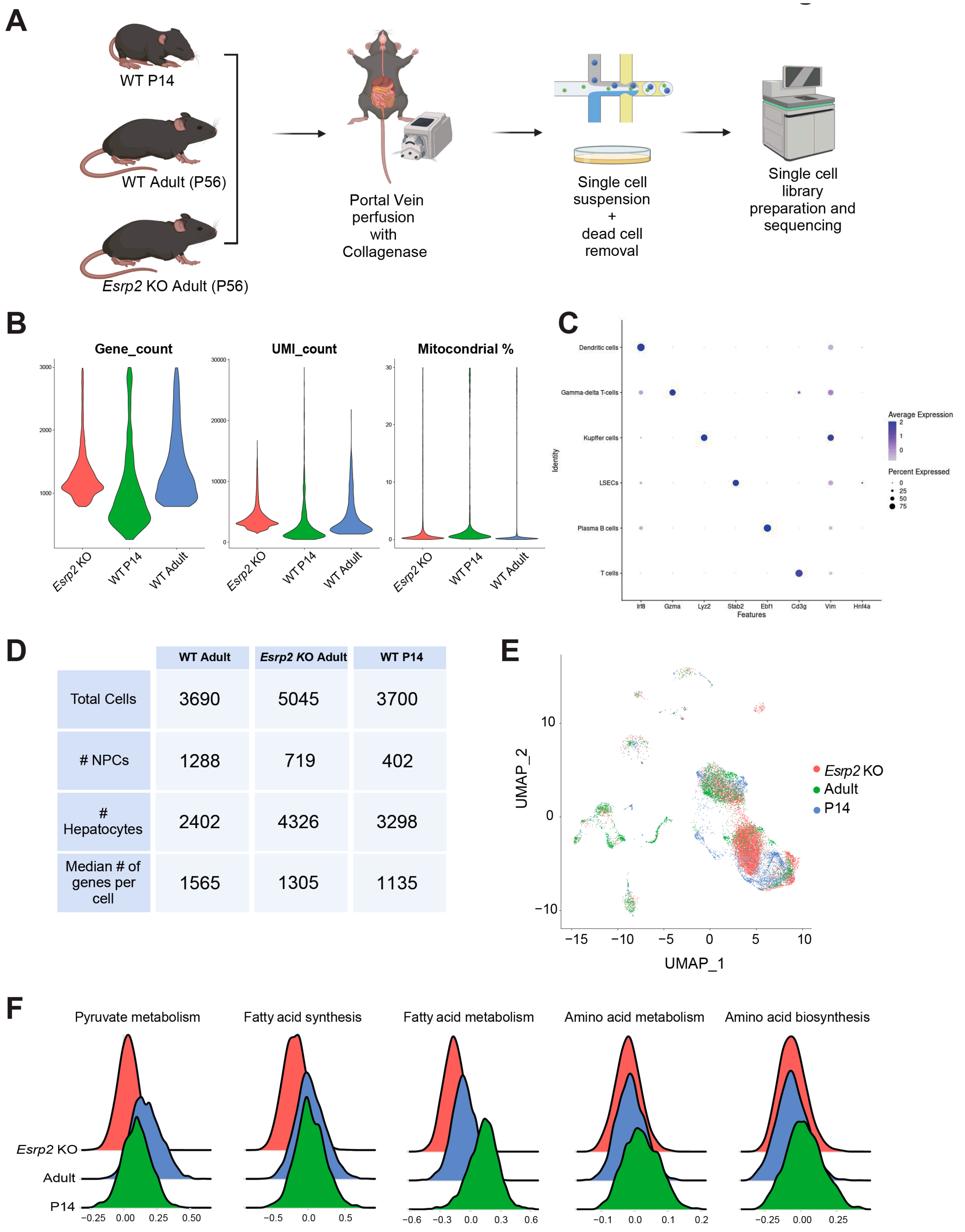


| | WT Adult | *Esrp2* KO Adult | WT P14 |
|---|---|---|---|
| Total Cells | 3690 | 5045 | 3700 |
| # NPCs | 1288 | 719 | 402 |
| # Hepatocytes | 2402 | 4326 | 3298 |
| Median # of genes per cell | 1565 | 1305 | 1135 |

**Supplementary Figure 5. (A)** Overview of scRNA-seq workflow for generating P14, adult, and *Esrp2* KO scRNA-seq libraries. Single cell library preparation was performed with whole cell suspensions individually for each mouse using 10X Chromium Single Cell 3' Reagent Kit (V3 chemistry) after magnetic-activated cell sorting to remove dead cells. (**B**) Violin plots showing the distribution of gene counts, UMI counts and % mitochondrial content in scRNA-seq libraries. (**C**) Dot plot showing the expression of cell type specific markers for NPCs from scRNA-seq libraries. (**D**) Table showing the numbers of cells and genes captured from the scRNA-seq libraries. (**E**) UMAP plot depicting the clusters of hepatocytes from P14, adult, and ESRP2KO scRNA-seq libraries. (**F**) Ridge plots of metabolic pathway scores for key adult metabolic functions in P14, adult, and *Esrp2* KO hepatocytes.

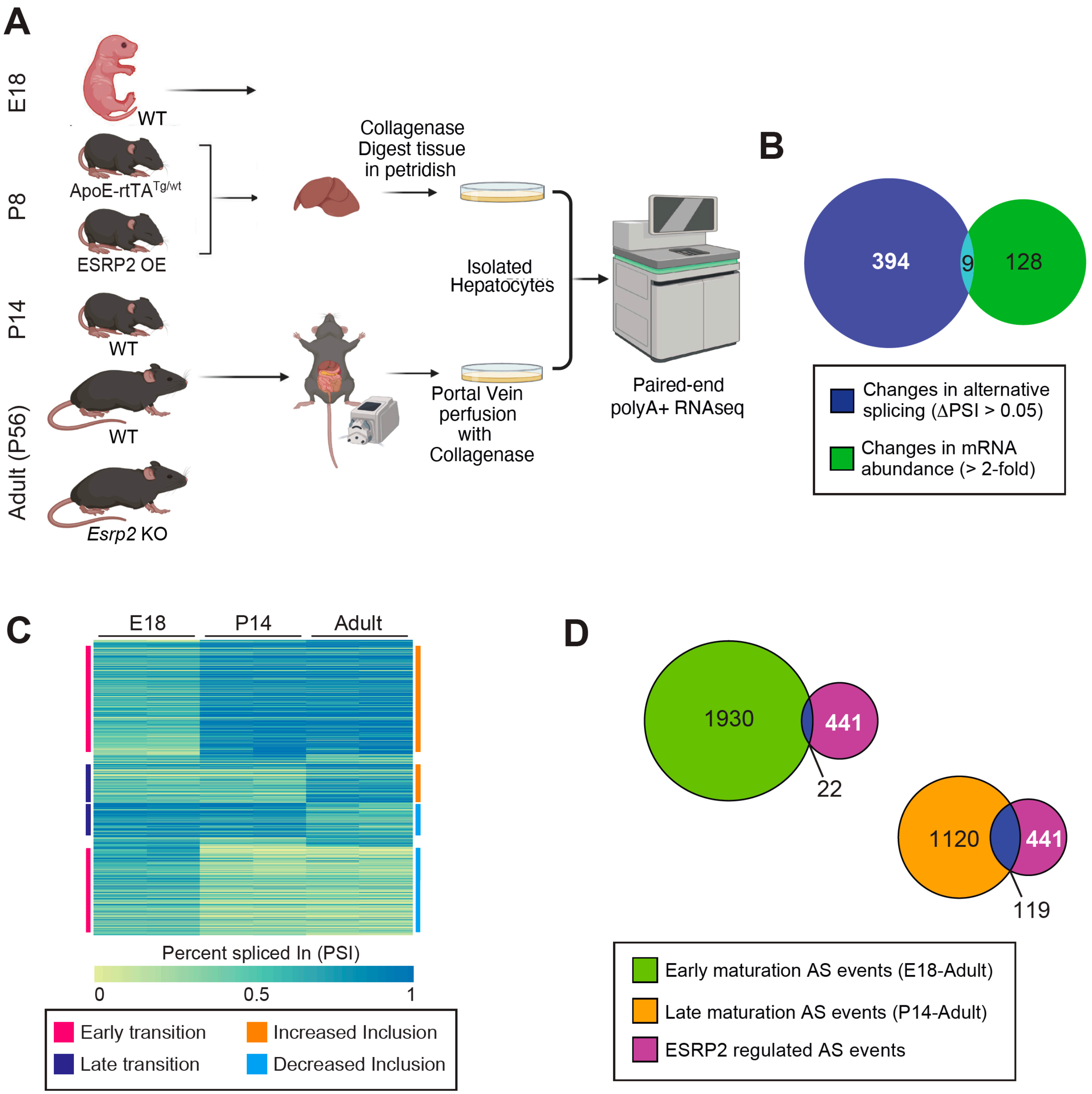


**Supplementary Figure 6. (A)** Overview of bulk hepatocyte-specific RNA-seq workflow for generating E18 WT, P8 Control and ESRP2 OE, P14 WT, and Adult WT and *Esrp2* KO. **(B)** Overlap of splicing and gene expression changes in WT vs. *Esrp2* KO Adult hepatocytes. **(C)** Heatmap showing alternative splicing transitions regulated across E18, P14 and Adult timepoints. PSI values across 3 conditions and 2 replicates were row normalized to plot the heatmap. **(D)** Overlap of alternative splicing (AS) events misregulated in *Esrp2* KO adult hepatocytes with Early (E18-P14) and Late (P14-Adult) postnatal transitions respectively.

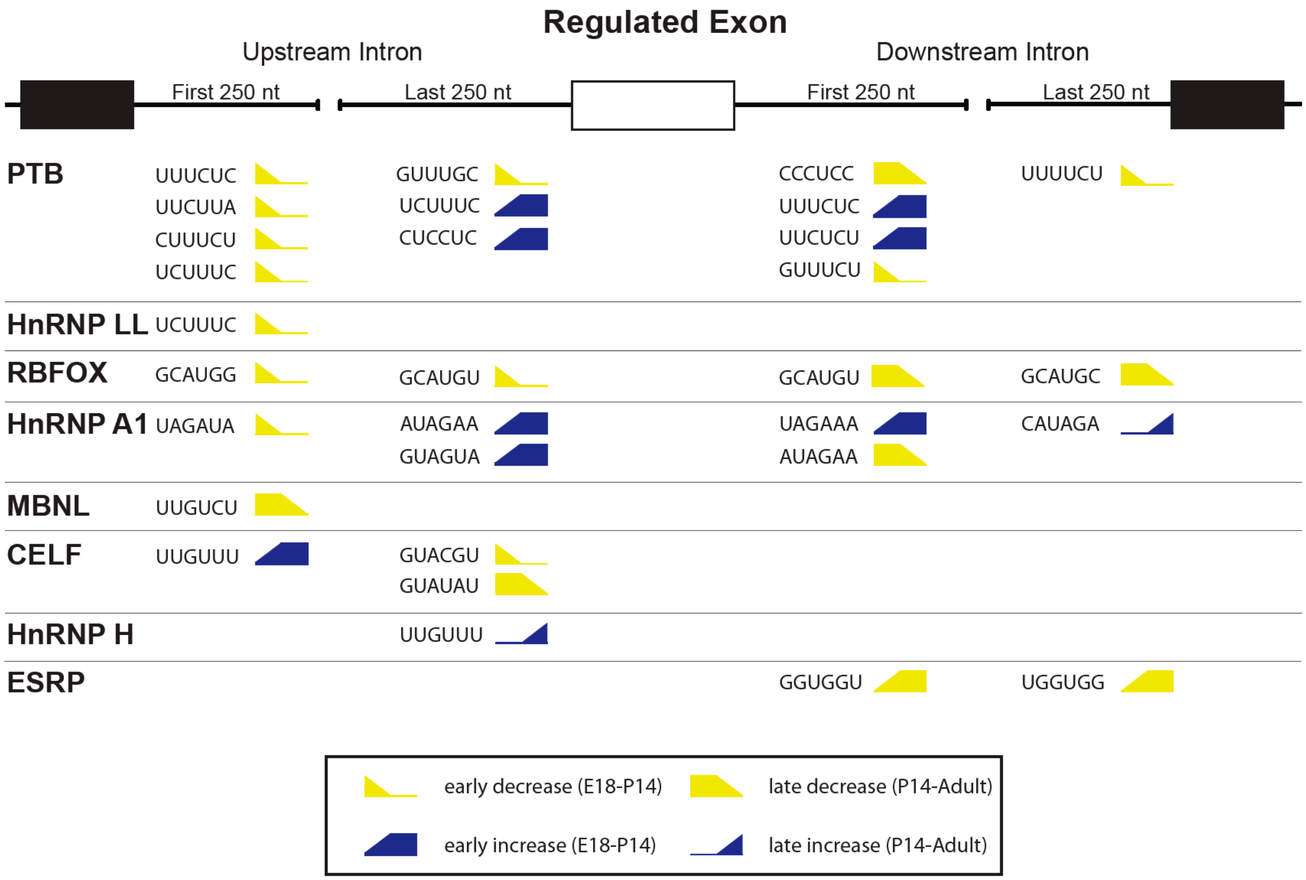


**Supplementary Figure 7.** Motif enrichment map of splicing factors around alternative exons regulated during postnatal liver development. Significantly enriched motifs (P<0.05) resembling known binding sites of splicing factors (in the four flanking 250-nt intronic regions) that also exhibit association with specific temporal transitions in the regression analysis are shown in yellow (decreasing inclusion) or blue (increasing inclusion) with icons indicating the transition pattern during postnatal liver development. ESRP (GGUGGU and UGGUGG) motifs were specifically associated with a late temporal pattern.

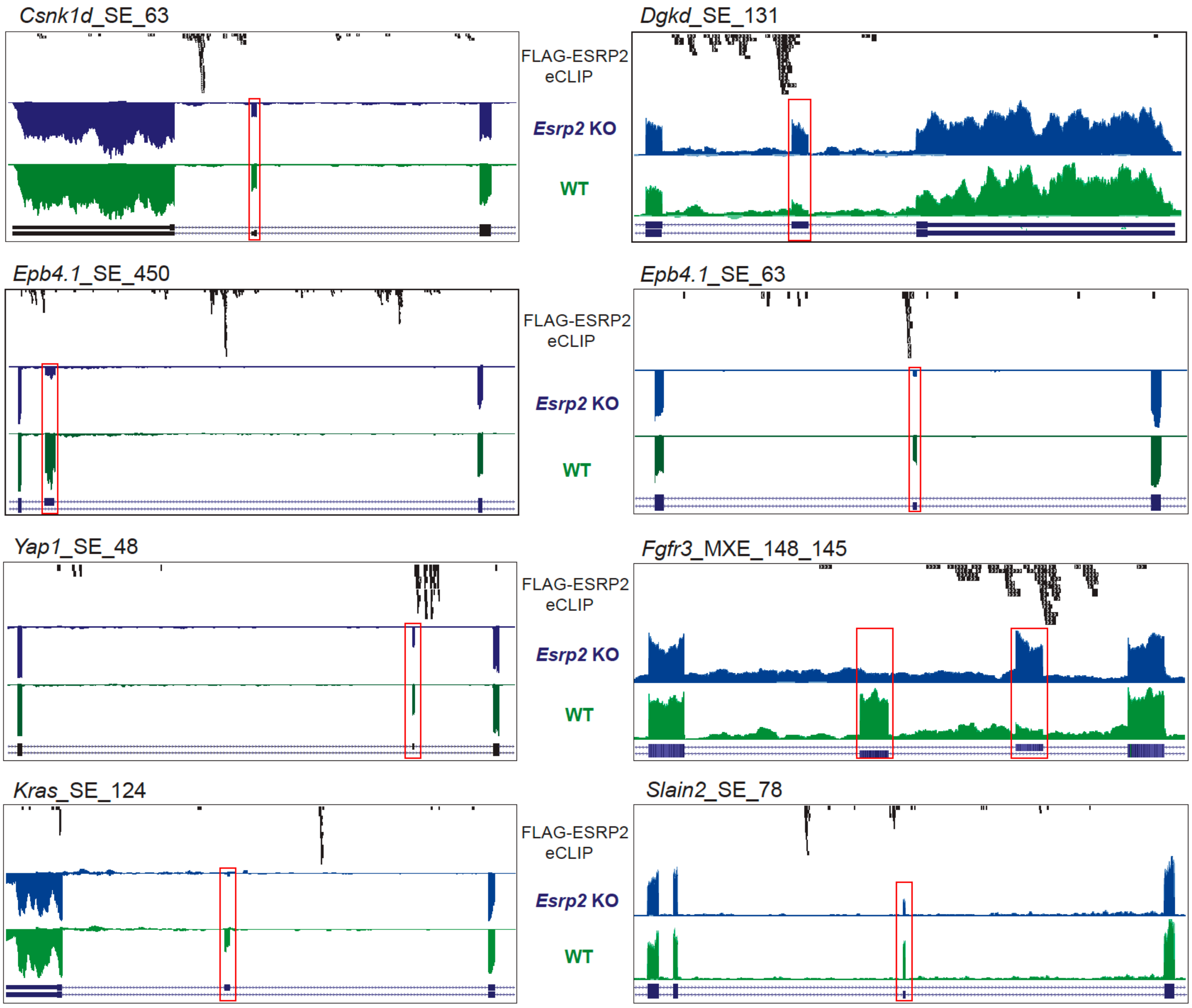


**Supplementary Figure 8.** Aligned genome browser tracks of direct ESRP2 target exons. RNA-seq tracks for eight representative alternative exons from WT (green) and *Esrp2* KO (blue) hepatocytes with ESRP2 eCLIP tags (black) in the surrounding intronic regions are shown. Boxes point to the regulated exons.

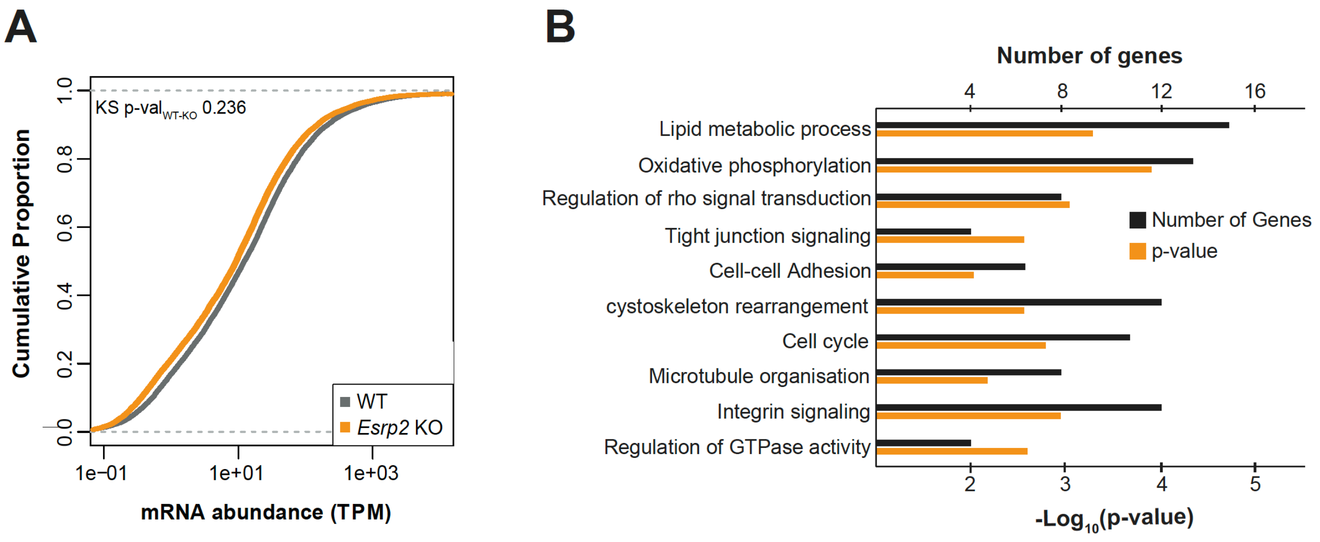


**Supplementary Figure 9. (A)** Cumulative plot of mRNA abundance for miR-122 non-regulated transcripts in RNA-seq data from WT and *Esrp2* KO hepatocytes. TPM; transcripts per million. p-value was calculated using Kolmogorov-Smirnov test. **(B)** Gene Ontology (GO) analysis of miR-122-regulated biological functions that are significantly affected in *Esrp2* KO mouse hepatocytes.
